# Specific targeting of MR1-antigen complexes using nanobodies

**DOI:** 10.64898/2026.03.22.713551

**Authors:** M.H. Shakir Hussain, Samuel J. Redmond, Wael Awad, Calvin Xu, Caroline Soliman, Lisa Ciacchi, Alexis P. Gonzalez, Jeffrey Y. W. Mak, David P. Fairlie, James McCluskey, Adam P. Uldrich, Jamie Rossjohn, Dale I. Godfrey, Hui-Fern Koay, Nicholas A. Gherardin

## Abstract

T cell receptor mimic (TCRm) antibodies and nanobodies that specifically bind peptide-HLA complexes have great therapeutic potential, as they can target polymorphic HLA on tumour cells furnishing peptides derived from tumour-associated antigens. MR1 is an MHC class-I-like molecule that exhibits limited polymorphism that binds and presents conserved metabolites, such as 5-OP-RU, derived from microbial riboflavin biosynthesis. Whether antibodies targeting such MR1-5-OP-RU complexes can be generated remains unclear. Using yeast display technology and *in vitro* affinity maturation, a nanobody with high affinity and fine specificity toward MR1-5-OP-RU complex was generated. These nanobodies bind both mouse and human MR1-5-OP-RU and inhibited MAIT cell responses to 5-OP-RU *in vitro* and *in vivo* demonstrating therapeutic potential. Moreover, we provide a molecular basis underpinning the fine specificity of these nanobodies, solving the crystal structures of MR1 in complex with either 5-OP-RU or Ac-6-FP. Here, the nanobody co-bound MR1 and 5-OP-RU, akin to a TCRm antibody. Moreover, we engineer bispecific antibodies targeting both MR1-5-OP-RU and CD3, that drive broad T cell killing of bacterially-infected cells as well as tumour cells treated with 5-OP-RU, thereby providing proof-of-principle for targeting the MR1 molecule with with TCRm-based nanobodies.

**One Sentence Summary:** We report the development of a nanobody targeting MR1-5-OP-RU complex and demonstrate its utility to modulate MAIT cells responses, and as a bispecific engager.

## Introduction

A key aspect of adaptive immunity is the ability of major histocompatibility complex (MHC) molecules to bind and present peptides at the cell surface(*1*). Antibodies, or nanobodies, targeting these peptide-Human Leucocyte Antigen (HLA) complexes, known as T cell receptor mimic (TCRm) antibodies, have gained clinical interest due to their ability to discriminate cells based on disease-associated peptides, such as those derived from tumour-associated antigens or pathogens(*2*). These TCRm antibodies are versatile, with applications that include engineering as full-length monoclonal antibodies (mAb), antibody-drug conjugates, or bispecific antibodies(*2, 3*). While promising, their therapeutic scope is nonetheless limited by the extreme polymorphism of HLA alleles, meaning that TCRm antibodies are only applicable to a subset of the human population(*4*). As a result, most TCRm platforms focus on HLA-A*02:01, the most prevalent HLA allele, yet its frequency varies widely across human populations(*5*).

MHC-related protein 1 (MR1) is an MHC I-like molecule that presents small metabolite-derived non-peptide antigens. Unlike the shallow, open binding cleft of MHC-I that binds short peptide antigens, the helical ‘jaws’ of MR1 features two closed pockets, one of which, the A¢ pocket, is lined with a series of aromatic residues enabling MR1 to capture and present small cyclic metabolites(*6*). These compounds frequently form a characteristic Schiff base covalent bond with a lysine residue, thereby tethering the ligands to the A¢ pocket(*6*). The most well-defined MR1 antigens are derived from microbial riboflavin biosynthesis whereupon the key biosynthetic intermediate, 5-amino-6-D-ribitylaminouracil (5-A-RU) reacts with small carbonyl compounds such as methylglyoxal to form intermediate pyrimidines like 5-(2-oxopropylideneamino)-6-D-ribitylaminouracil (5-OP-RU)(*7*). Here, the carbonyl moiety froms the carbonyl compounds form a covalent bond with MR1, and the ribityl chain of 5-A-RU projects up toward the opening of the A¢ pocket, poised for TCR immunosurveillance by an evolutionary conserved T cell lineage known as mucosal-associated invariant T (MAIT) cells(*7*). Given the evolutionary conservation of riboflavin synthesis in many microbes, 5-A-RU adducts serve as a molecular signature of infection. Also, in contrast to MHC-I which is constitutively expressed at high levels on the surface of nucleated cells, MR1 is largely retained intracellularly in the endoplasmic reticulum (ER) with only low cell surface expression(*8*). However, MR1 rapidly traffics from the ER to the surface at higher levels upon ligand engagement(*9*). Importantly, MR1 has very little variation between indivuals, other than some rare allelic variants(*10*), making MR1-antigen complexes attractive population-wide therapeutic targets for novel TCRm antibodies that may, for example, block pathological MR1-antigen driven MAIT cell responses. Alternatively, MR1-antigen complexes on the surface of tumour cells or infected cells may represent targets for TCRm antibody-based therapeutics.

Targeting MR1-antigen complexes with antibodies or nanobodies poses unique challenges however. The presented antigens are small and typically buried within the A¢ pocket, leaving only a minimal epitope exposed for TCR recognition(*11*). Indeed, structural studies show that when the MAIT TCR docks on MR1-5-OP-RU complexes, the only conserved interaction between the TCR and the 5-OP-RU antigen is a hydrogen bond(*12*). Thus, the discovery of antibodies with fine specificity toward small motifs within a large, conserved protein molecule using traditional animal immunisation protocols is challenging, often involving large, high-throughput screening approaches(*13*). To overcome these challenges, *In vitro* display systems, such as yeast surface display, however, can overcome this challenge, offering a more refined approach that allows rapid positive and negative selection of clones that can distinguish between highly similar proteins(*14*). Moreover, the single domain of nanobodies (also known as single-domain antibodies or variable domain of heavy chain antibodies; VHH), derived from the camelid family of mammals, are inherently suited to gripping crevices and to binding small, buried epitopes due to their compact, single-domain structure(*15*). Indeed, yeast display of nanobodies has been employed successfully to isolate binders with fine molecular specificity, for example, nanobodies that distinguish conformation-specific species of G-protein coupled receptors (GPCRs)(*16*).

Here, we devised a platform approach for the generation of antigen-specific MR1-binding nanobodies, incorporating an established synthetic nanobody yeast-display library(*14*). We used this platform to generate nanobodies with fine specificity toward MR1-5-OP-RU complexes, and identified a lead candidate, clone C11, with sub–nanomolar affinity and selectivity for 5-OP-RU bound to both mouse and human MR1. We show that C11 can modulate antigen-driven MAIT cell activity *in vitro* and *in vivo*, and provide a molecular basis for this specificity towards MR1-5-OP-RU. Moreover, we engineer C11 into bispecific T cell engagers that selectively redirect T cells to bacteria-infected antigen-presenting cells and to tumour cells treated with synthetic 5-OP-RU. These findings pave the way for the therapeutic targeting MR1-antigen complexes, offering a new approach to the application of TCRm antibodies.

## Results

### Nanobody selection and enrichment identifies MR1-antigen binders

To generate nanobodies with fine-specificity toward MR1-antigen complexes, a synthetic nanobody yeast display library(*14*) was used, together with a selection strategy incorporating sequential negative and positive enrichment steps. This iterative enrichment approach combines magnetic (MACS) and flow cytometric (FACS) selection of yeast prior to cloning, first depleting the yeast library of binders with unwanted specificity, followed by enrichment with probes of desired specificity, providing an efficient and rapid method for isolating high-specificity nanobodies, in as quickly as two weeks (**Figure 1A**)(*14*). Here, initial rounds involved anti-PE magnetic bead-based MACS-depletion of non-specific binders from the naïve yeast library with PE-Cy7–labelled human (hu)CD1c tetramers, removing clones that recognise streptavidin, β2-microglobulin, PE-fluorophore conjugates, or the magnetic beads. The flow-through was then positively selected using PE-labelled huMR1–5-OP-RU tetramers, and placed into culture to expand selected cells. After three rounds of similar MACS selection, yeast libraries were sorted using FACS, where nanobody expression at the surface of yeast was detected via a co-expressed hemagglutinin (HA) tag, enabling identification of HA^+^ yeast displaying surface nanobody. FACS sorting depleted PE-Cy7-labelled human (hu)CD1c reactive cells and purified single positive cells PE-labelled huMR1–5-OP-RU tetramers that bound huMR1–5-OP-RU tetramer (**Figure 1A & B**). Reanalysis of these FACS-sorted yeast after culture demonstrated clear enrichment relative to the naïve library for yeast that bound huMR1-5-OP-RU tetramers but not huCD1c tetramers (**Figure 1B**). A substantial proportion of these also bound huMR1 tetramers loaded with 6-formylpterin (6-FP), indicating that many clones recognised MR1 in an antigen-independent manner. However, a higher proportion bound to huMR1-5-OP-RU (51.6%) over huMR1-6-FP (38.8%), suggesting ligand discrimination by some clones. Approximately half of the enriched clones also bound mouse (m)MR1-5-OP-RU (22.6%), reflecting the high sequence conservation between the a1 and a2 domains of human and mouse MR1, particularly around amino acid side-chains surrounding the ligand-binding A¢ pocket (**Figure 1B, S1**). These data indicated successful enrichment of clones with 5-OP-RU–dependent and cross-species MR1 specificity, prompting single cell sorting to generate clonal colonies of yeast that only bound the huMR1-5-OP-RU tetramers (**Figure 1C**).

**Fig. 1.**
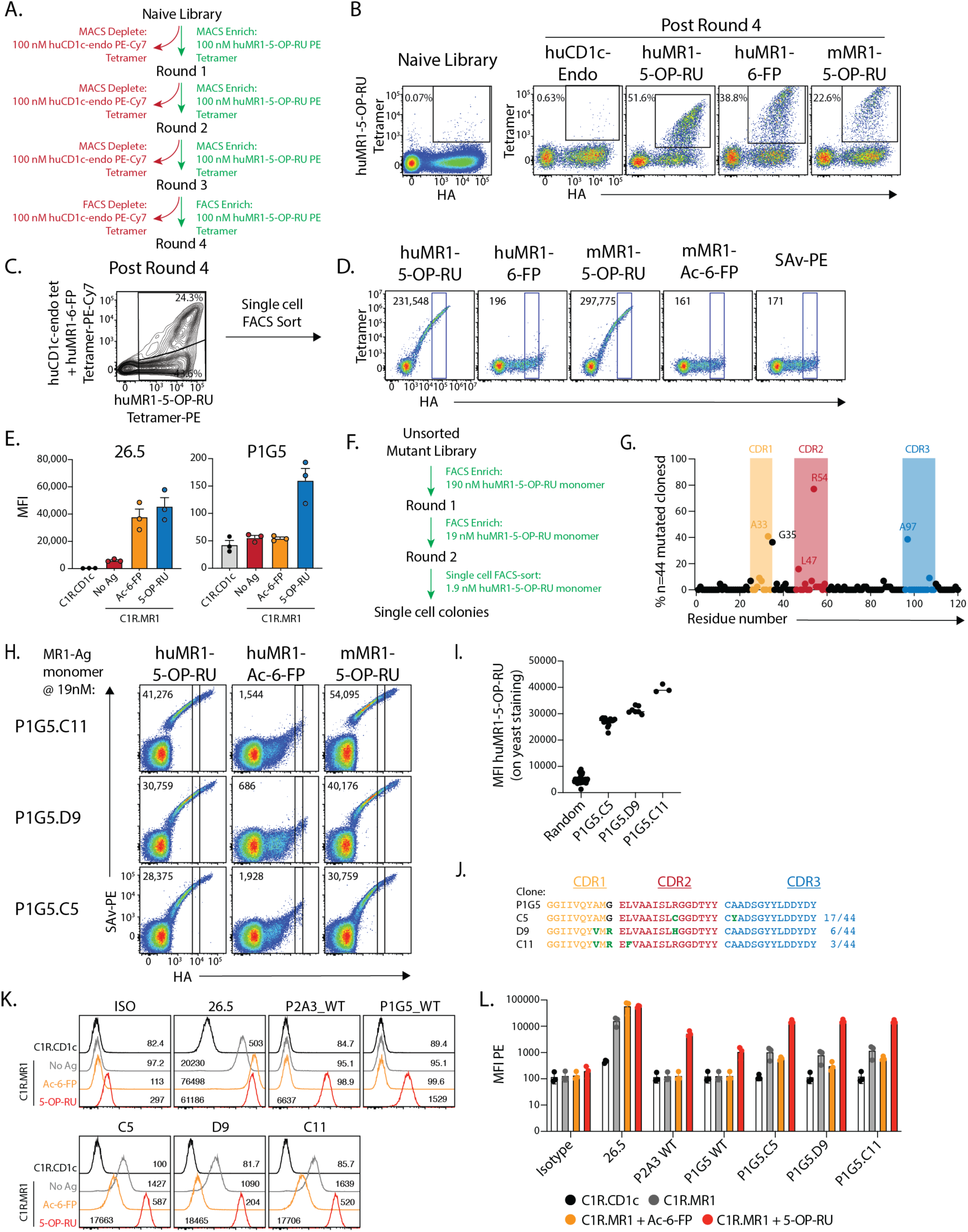
Development of an MR1-antigen-specific nanobody. **A.** Schematic demonstrating the selection strategy for the first four rounds of yeast selection. **B.** Flow cytometric pseudocolour plots showing tetramer staining profiles on the yeast library after four rounds of selection compared to the naive library. hu, human; m, mouse. **C.** Flow cytometric contour plot showing co-staining with PE or PE-Cy7-conjugated tetramers on HA^+^ yeast after four rounds of selection. **D.** Flow cytometric pseudocolour plots showing MR1-antigen tetramer staining on a single-cell sorted colony, clone P1G5. Numbers indicate median fluorescence intensity (MFI) of the gated population. **E.** Bar graph showing MFI of anti-MR1 clone 26.5 antibody or P1G5-Fc on C1R cells treated with Ac-6-FP or 5-OP-RU. Bars show the mean of n = 2 technical replicates from each of n = 3 independent experiments. **F.** Schematic demonstrating yeast selection strategy for affinity maturation yeast libraries. **G.** Scatter graph showing frequency of mutations for individual amino acids of the P1G5 nanobody. Each datapoint represents an individual amino acid and the frequency corresponds to how often that amino acid was mutated across the 44 sequenced clones. **H.** Representative flow cytometry pseudocolour plots showing tetramer staining after single cell sorting on yeast colonies with sequences aligning with P1G5 subclones C11, D9 and C5. Numbers indicate MFI of the gated population. **I.** MFI of huMR1-5-OP-RU monomer staining gated as per H for all colonies that had C5, D9 or C11 subclone sequences. Colonies with all other sequences are grouped in the ‘random’ column. **J.** Alignment of CDR amino acid sequences highlighting mutations between P1G5 and its subclones. **K**. Flow cytometric histogram overlays showing anti-MR1 antibody clone 26.5 or Nb-Fc staining on C1R cells after treatment with Ac-6-FP or 5-OP-RU. **L.** MFIs derived from K depicting n=3 independent experiments. Numbers on plots in D and H correspond to MFI of gated population whereas numbers in K correspond to MFI of total population. Error bars in E and L are standard error of the mean (SEM).

Two hundred yeast clones were isolated and screened for binding to a panel of PE-labelled tetramers including: huMR1-5-OP-RU, huMR1-6-FP, mMR1-5-OP-RU, mMR1-Ac-6-FP and streptavidin-PE alone. Clones exhibited diverse staining profiles including: (i) binding huMR1-5-OP-RU but not huMR1-6-FP, with no mouse MR1 cross-reactivity; (ii) clones exclusively binding huMR1 regardless of antigen; (iii) clones binding all MR1 variants irrespective of antigen and species (**Figure S2A**) irrespective of species or antigen; and (iv) the desired clones with exclusive specificity for MR1-5-OP-RU across both human and mouse (**Figure 1D, S2A**). No clones bound streptavidin-PE alone. Sequencing of desired clones (archetypal clones shown in **Figure S2B**) revealed seven unique nanobody sequences with high CDR loop sequence variation (**Figure S2C**). These nanobody clones were next expressed as mouse IgG1 Fc-fusion dimers (Nb-Fc) and tested for their ability to stain a control CD1c-expressing C1R cell line (C1R.CD1c) or MR1-overexpressing C1R cells (C1R.MR1) treated with Ac-6-FP or 5-OP-RU (**Figure 1E & Figure S2D**). The reference anti-MR1 antibody clone 26.5, which binds MR1 independently of antigen, stained C1R.MR1 cells to a similar degree following Ac-6-FP or 5-OP-RU treatment, in line with ligand-induced MR1 surface upregulation(*8*). In contrast, only three Nb-Fc clones, P1G5, P2A3 and P2E8, bound MR1 in a 5-OP-RU-dependent manner. Their overall staining intensity was weak relative to 26.5, suggesting low-affinity binding, despite high intensity on-yeast staining.

### In vitro affinity maturation generates high-affinity MR1-Antigen binders

The two nanobody clones that labelled brighter for MR1-5-OP-RU, namely P1G5 and P2A3, were next selected for *in vitro* affinity maturation using error-prone PCR, resulting in new mutant yeast-nanobody libraries generated for each clone (**Figure S3A**). To enhance resolution of binding differences, mutant libraries were screened using huMR1-5-OP-RU monomer rather than tetramer, resulting in a range of staining intensities likely due to reduced avidity (**Figure S3B**). This monomer-based staining approach enabled differentiation and selection of higher-affinity mutants across three rounds of FACS enrichment for yeast with the highest staining intensities, with a 10-fold reduction in monomer concentration also applied across rounds to further favour high-affinity binders. By the final round of single cell sorting to grow clonal colonies as was performed in the initial selection strategy (**Figure 1F**), the libraries were highly enriched for clones retaining MR1-5-OP-RU binding while excluding reactivity to MR1-Ac-6-FP (**Figure S3C**). Forty-four P1G5 and 72 P2A3 single-cell subclones were screened, each of which retained strong cross-species MR1-5-OP-RU binding and minimal huMR1-Ac-6-FP binding.

Sequencing of these clones revealed a series of overlapping mutations largely within the CDR loops (**Figure 1G, S3E**). Three prominent P1G5-derived subclones C5, D9 and C11, accounted for the highest intensity huMR1-5-OP-RU staining with some overlap between mutations (**Figure 1I & J**), while the top P2A3 mutant, F2, bore a frequently occuring Tyr-to-Cys substitution at CDR3H position 95, but was poorly expressed (**Figures S3E & F**). Accordingly, Nb-Fc expression and functional analysis focused on the top P1G5 derivatives, where C5, D9, and C11 Nb-Fc dimers showed improved staining of 5-OP-RU-pulsed C1R.MR1 cells relative to their parental ‘wildtype’ clone (C1R.wt) and control antibodies. (**Figure 1K & L**). Although some binding to MR1-Ac-6-FP, and MR1 on unpulsed cells, was detectable, especially in clones with higher avidity, clone C11 stood out with the strongest 5-OP-RU signal and relatively preserved ligand selectively. Ultimately, D9 and C11 emerged as lead clones achieving high staining intensity for 5-OP-RU pulsed cells, with preserved antigen specificity, and were prioritised for downstream characterisation.

### Biochemical characterisation of MR1-antigen specific nanobodies

To help define the molecular basis of MR1-antigen binding by these nanobodies, mutational mapping was first performed using a panel of C1R cell lines expressing MR1 with single point mutations along the a1 and a2 helices(*17*). C1R cells were treated with 5-OP-RU and stained with either C11-Fc or reference mAb 26.5 (**Figure 2A**). Staining by C11-Fc was impaired by R61A and L65A mutations, both adjacent to the A¢ pocket on the a1 helix, consistent with proximity to the antigen-binding cleft (**Figure 2A**). In contrast, staining with 26.5 mAb was reduced by N146A and H148A mutants on the opposite side of the antigen-binding cleft along the a2 helices, closer to the F¢ pocket (**Figure 2A**). These non-overlapping profiles suggest distinct binding modes. To formally assess binding affinities between the nanobody and MR1, surface plasmon resonance (SPR) was performed using immobilised MR1 loaded with defined ligands (**Figure 2B**). The parental wild-type P1G5 nanobody bound huMR1-5-OP-RU with an affinity constant (KD) of ∼ 3.2 mM, while the affinity-matured C11 mutant exhibited ∼ 1500-fold improved affinity, with a KD of ∼ 1.8 nM (**Figure 2B**). Both nanobodies showed similar degrees of reactivity to mMR1-5-OP-RU with similar binding affinities (KD ∼ 2.9 mM and 0.5 nM for the wild type and C11 respectively, **Figure 2B**). While the parental nanobody failed to bind huMR1-Ac-6-FP in the concentration range measured, C11 showed a weak interaction of KD ∼ 20 mM, over 10,000-fold weaker than that for huMR1-5-OP-RU. While 26.5 mAb bound both huMR1-5-OP-RU and huMR1-Ac-6-FP with comparable affinity constants of KD ∼ 5.1 nM and 2.1 nM respectively, consistent with a lack of specificity for particular MR1-bound antigens. Notably, 26.5 mAb bound huMR1-5-OP-RU with ∼ 3-fold lower affinity than C11 (KD ∼ 1.7 nM), while binding to mMR1-5-OP-RU was comparable between the two (**Figure 2B**). Notably, C11 has faster on-rates and slower off-rates toward huMR1-5-OP-RU and mMR1-5-OP-RU compared to 26.5 mAb as well as the wild-type P1G5 nanobody (**Figure 2B**). Collectively, these results highlight the marked effect of *in vitro* affinity maturation on the strength of nanobody binding, and demonstrate the antigen-specificity of the C11 nanobody for MR1-5-OP-RU.

**Fig. 2.**
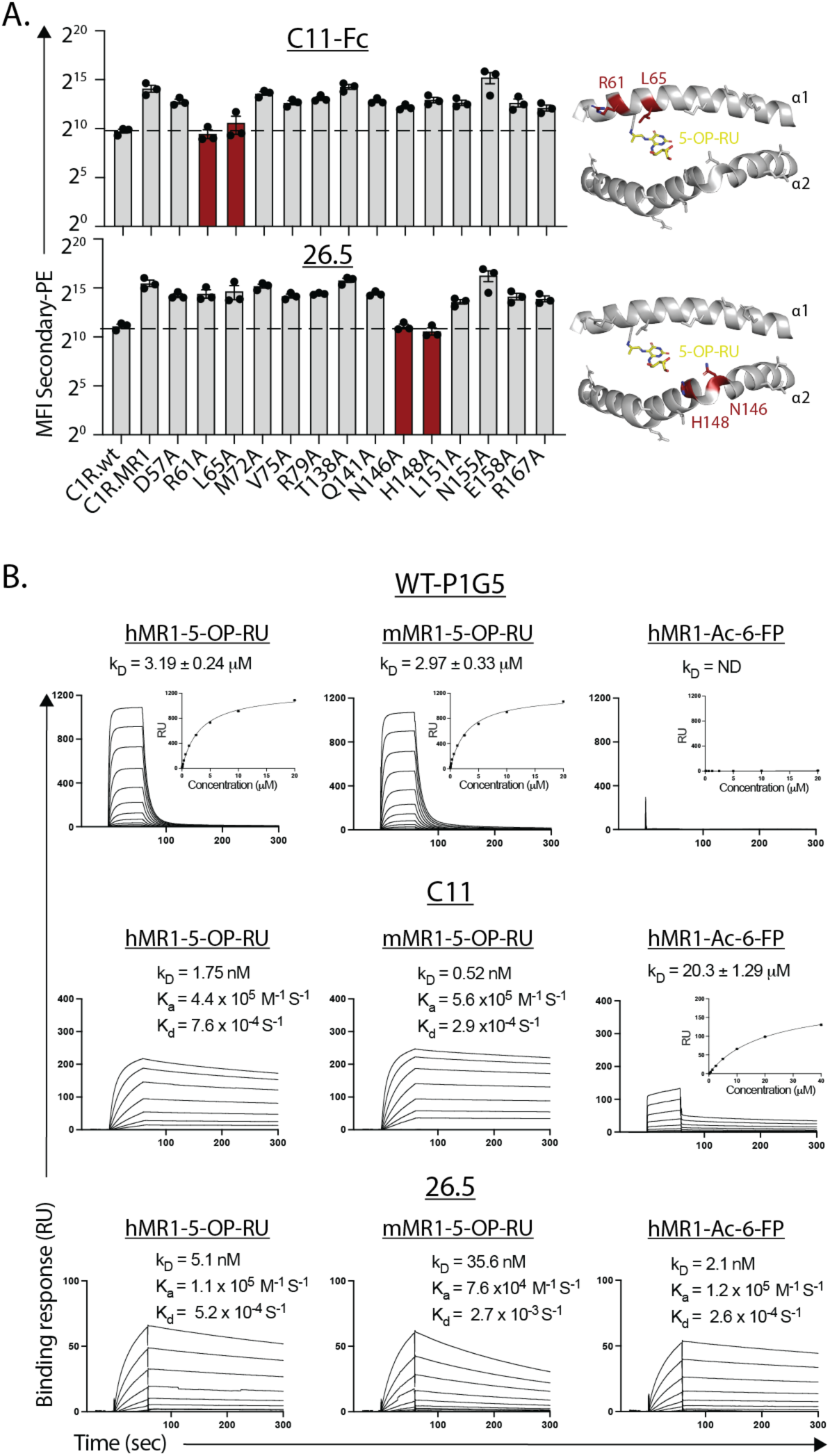
Biochemical characterisation of C11 binding to MR1-antigen complexes. **A.** Left: Bar graph showing median fluorescence intensity (MFI) of PE-conjugated secondary anti-mIgG antibody staining of C1R cell lines overexpressing MR1 or alanine point mutants, after pulsing with 5-OP-RU and staining with either C11-Fc or 26.5 antibody. Dotted line is a visual representation of the MFI for C1R.wt staining. Red bars highlight mutant cell lines reduced to C1R.wt levels. Data points represent mean of n=2 technical replicates from n=3 independent experiments. Error bars depict standard error of the mean. Right: Cartoon representation of MR1 antigen-binding cleft (PDB 5D5M) highlighting position of mutations with mutants affecting binding highlighted in red. **B.** Surface plasmon resonance sensorgrams showing steady-state binding affinities of hMR1-5-OP-RU, mMR1-5-OP-RU and hMR1-Ac-6-FP for WT nanobody (*upper panel*) as well as kinetics or steady state binding of titrated concentrations of hMR1-5-OP-RU, mMR1-5-OP-RU and hMR1-Ac-6-FP to immobilized C11 nanobody (*middle panel*) and 26.5 antibody (*lower panel*). Data representative of n = 2 technical replicates. The SPR sensograms and equilibrium curves (shown as insert panel in the steady state affinity runs) were prepared in GraphPad Prism 10. Error bars in the equilibrium curves represent the mean and SD from technical replicates. ND, not determined. RU, response unit.

### The molecular basis underlying C11 nanobody binding of MR1-antigen complexes

To understand the structural foundation of nanobody antigen-specificity, the crystal structure of C11 in complex with huMR1-5-OP-RU and huMR1-Ac-6-FP were determined at resolutions of 2.9Å and 3.1Å respectively (**Figure 3 and Table S1**). The electron densities of the 5-OP-RU and Ac-6-FP ligands and the interfaces between the C11 and MR1-antigen complexes were clear. The C11 molecule docked atop the antigen binding cleft of MR1-5-OP-RU and MR1-Ac-6-FP in a manner analogous to the docking of MAIT TCR on MR1 (**Figure 3A-F**). Despite its relatively small size, the C11 nanobody, when interacting with MR1-5-OP-RU, exhibited a comparable buried surface area (BSA) when bound to MR1, with a value of 1165 Å^2^, which is similar to that of the MAIT TCR-MR1-antigen complex (BSA of 1170 Å^2^). However, in the crystal structure of the C11-MR1-Ac-6-FP complex, the C11 displayed conformational alterations in the CDR1H loops (discussed below), resulting in a significantly reduced BSA upon complexation of 830 Å^2^ (**Fig. 2B**).

**Fig. 3.**
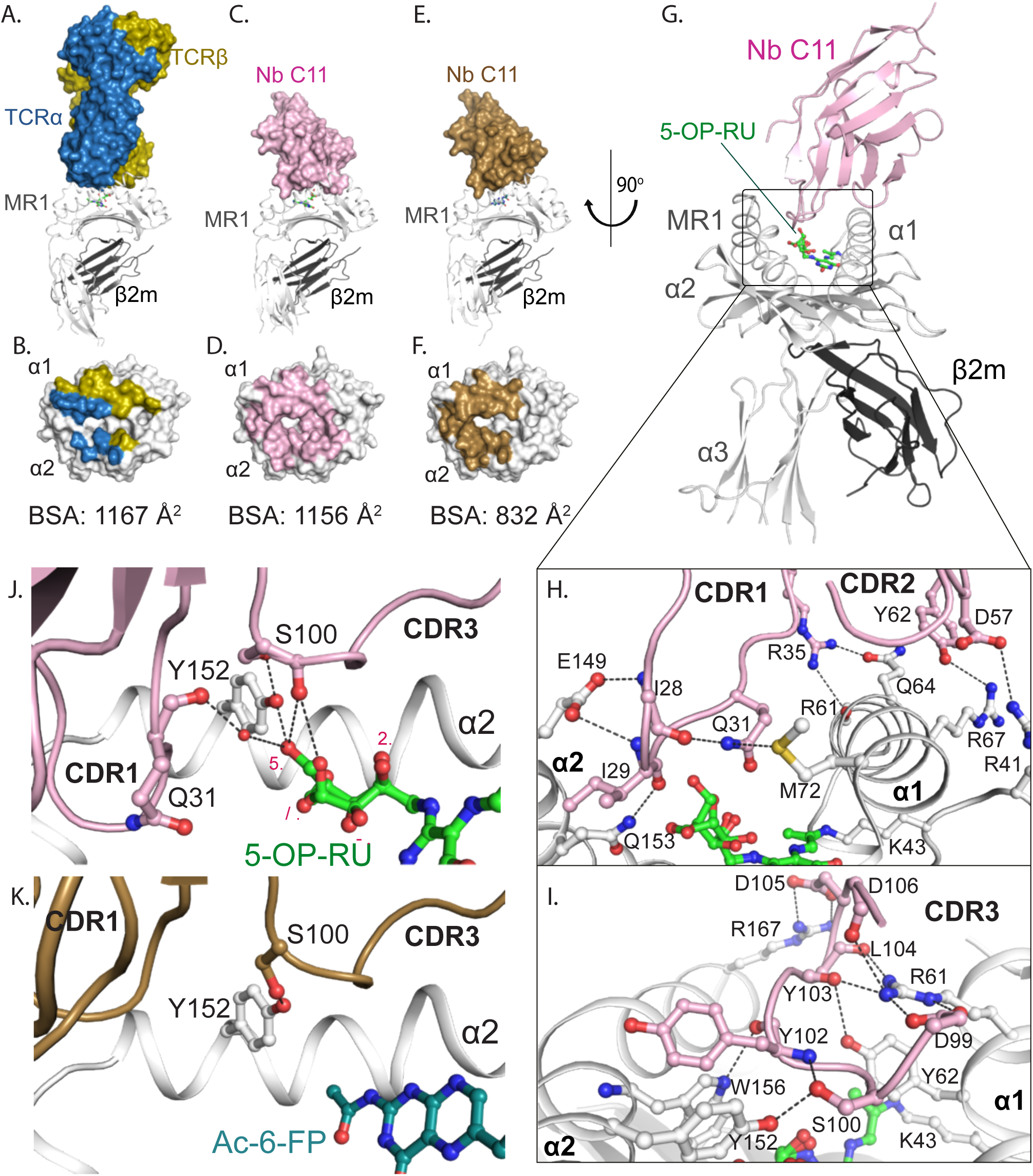
The molecular basis for P1G5.C11 binding to MR1-5-OP-RU and MR1-Ac-6-FP. **A-F** Crystal structures of (**A-B)** the ternary complex of MAIT A-F7 TCR-MR1-5-OP-RU (PDB; 6PUC), (**C-D**) the C11-MR1-5-OP-RU complex, and (E-F) C11-MR1-Ac-6-FP complex. The top panels (**A, C and E**) are representations of the complexes; the lower panels (**B, D and F**) illustrate the respective (**B**) TCR and (**D and F**) C11 footprints on the molecular surface of MR1-antigen complexes. The MR1 and b2-microglobulin molecules are colored white and dark grey, respectively. A-F7 TCRa, sky-blue; TCRb, olive; C11 in complex with MR1-5-OP-RU, pink; and C11 in complex with MR1-Ac-6-FP, sand. 5-OP-RU and Ac-6-FP are shown as green and sky-blue sticks, respectively. **G.** Cartoon representation of the structure of C11-MR1-5-OP-RU. **H-I.** Interactions between (**H**) the CDR1H and CDR2H loops, (**I**) CDR3H loop of the C11 and MR1-5-OP-RU residues. The interacting residues are shown as sticks. The hydrogen bonds interactions are represented as black dashes. **J-K**. Interactions of the CDR1 and CDR3 of the C11 nanobody in complex with (**J**) MR1-5-OP-RU, and (**K**) MR1-Ac-6-FP, depicting the interactions between CDRH loops, MR1-residues and the binding ligands.

In the C11-MR1-5-OP-RU structure, most interactions were primarily associated with the CDR3H loop, while the CDR1H and CDR2H loops were involved to a significantly lesser extent (**Figure 3G-I**). In this context, the CDR1H loop was extended over the antigen binding cleft of MR1 and deeply embedded within the A¢ pocket, sandwiched between the a1 and a2 helices of MR1. Specifically, the backbones of the CDR1H-Ile28 and Ile29 residues established H-bonds with the side chains of Glu149 and Gln153 from the a2 helix of MR1, respectively. On the other side, the side chains of CDR1H-Gln31 and Arg35 formed H-bonds with MR1-Met72, Arg61 and Gln67 from the a1 helix of MR1. Here, CDR1H-Val33 formed a hydrophobic interactions with MR1-Leu65, that explained the impact of the Leu65Ala mutation on the C11 binding. Furthermore, the CDR2H loop of C11 was situated atop the a1 helix, creating two H-bonds between CDR2H-Asp57 and Tyr62 with MR1-Arg41 and Arg67 respectively. Notably, the CDR3H loop wedged between the helical jaws of the MR1 cleft, surrounding the F¢ pocket, although proximal to the bound antigen in the A¢ pocket. Here, the “^99^Asp-Ser-Gly-Tyr-Tyr-Leu-Asp-Asp^106^” segment plays a crucial role in mediating CDR3H interactions with MR1 (**Figure 3I**). The backbone of this peptide folded upon itself, effectively capping the F¢ pocket and engaging extensively with various residues of the MR1 α1- and α2-helices. Specifically, CDR3H-Asp99, Tyr103, Leu104 and Asp106 established an extensive network of H-bonds and van der Waals interactions with the MR1-Arg61 residue located in the a1 helix. This explains the impact of Arg61Ala mutation on C11 recognition of MR1 (**Figure 2A**). On the other side, the CDR3H-Ser100, Tyr102, Asp105 residues formed H-bonds with the MR1-Tyr152, Trp156 and Arg167 residues from the α2 helix, respectively. Accordingly, the CDR3H loop plays a prominent role in the interaction and recognition of the C11 nanobody by the MR1-5-OP-RU complex.

The conserved Tyr95a in CDR3a of the MAIT TCR is the responsible for trigerring MAIT cell activation by forming a network of H-bond interactions with 5-OP-RU and MR1-Tyr152(*18*). Here, we found that the CDR3H-Ser100 residue is oriented towards the bound antigen (5-OP-RU) within the A¢ cleft, establishing a direct H-bond with the 5¢-OH of one ribityl conformation, as well as a hydrogen bond with the 4¢-OH of the other ribityl conformation (**Figure 3J**). Additionally, the carbonyl group of C11-Gln31 formed a water-based H-bond with the 5′-OH of one ribityl conformation. Notably, the crystal structure of C11-MR1-Ac-6-FP revealed that the CDR1H loop was flipped out of the pocket and the CDR3H loop exhibited no direct contacts with Ac-6-FP (**Figure. 3K**).

Collectively, C11 nanobody established extensive hydrophilic and hydrophobic interactions with MR1-5-OP-RU, compared to much less buried surface and contacts with MR1-Ac-6-FP. This elucidates the strong binding affinity of C11 for MR1-5-OP-RU, whereas a weaker interaction was observed with MR1-Ac-6-FP.

### Functional blockade of MAIT cell activation by MR1-5-OP-RU specific nanobodies

To test the bioactivity of C11 and D9-Fc, they were assessed in a series of *in vitro* assays using cell lines and primary MAIT cells. First, a reporter SKW-3 cell line expressing a canonical MAIT TCR, clone MBV28(*19*), (SKW-3.MAIT) was co-cultured with C1R.MR1 cells in the presence of graded quantities of 5-OP-RU (**Figure 4A**). Both C11 and D9-Fc blocked SKW-3.MAIT cell activation as measured by CD69 upregulation, to a similar degree as clone 26.5 which is the gold-standard antibody for blocking MR1-dependent T cell activation. As anticipated, the isotype (ISO; another Fc-linked nanobody) and ‘no block’ controls did not inhibit activation, with similarly high MFI values for CD69 in response to all doses of 5-OP-RU. Blocking was 5-OP-RU-dependent, as C11 had no effect on SKW-3 cell lines that respond to endogenous MR1-presented ligands, unlike clone 26.5 which broadly inhibited all activation. Specifically, C11 blocked 5-OP-RU–driven activation of the MBV28 MAIT clone, but not ligand-independent responses from clones DGB129(*20*) or MC.7.G5(*21*) (**Figure S4**).

**Fig. 4.**
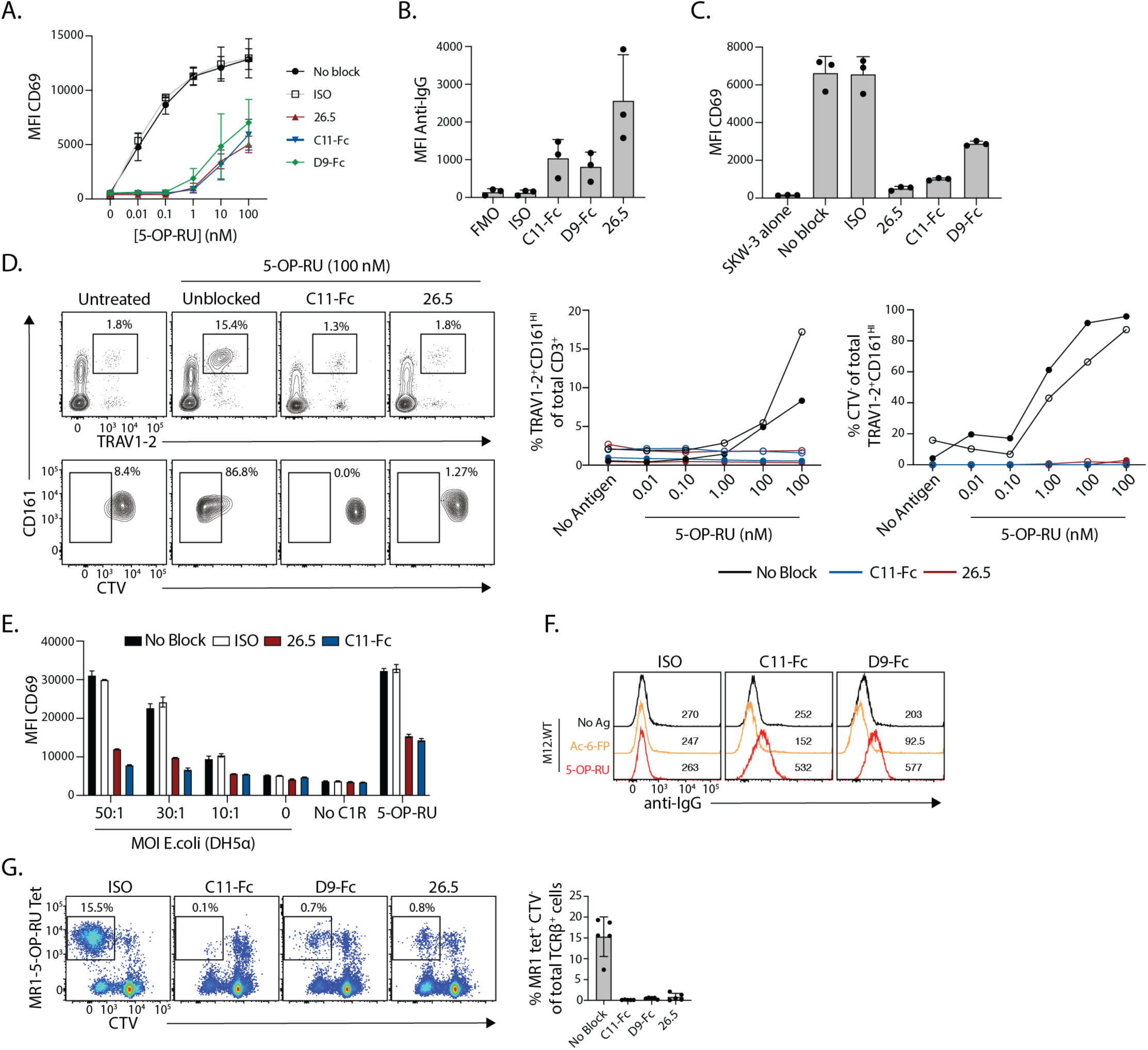
Characterisation of nanobody specificity and bioactivity. **A.** Line graph showing CD69 expression on SKW-3.MAIT cells after overnight culture with C1R.MR1 cells and 5-OP-RU in the presence or absence of nanobodies or controls. **B.** Bar graph showing staining intensity of nanobodies on C1R.wt cells after 3 hr pulse with 10 mM 5-OP-RU. **C.** Bar graph showing CD69 expression on SKW-3.MAIT cells after co-culture with C1R.wt cells plus 5-OP-RU. Data points in A-C are mean of n=2 technical replicates, showing n=3 independent experiments. **D.** Left: Representative flow cytometric contour plots gated on total CD3^+^ T cells (*top*) or TRAV1-2^+^CD161^HI^ MAIT cells (*bottom*) showing blocking of MAIT cell expansion and CTV dilution within PBMC after 6-day *in vitro* culture with 100 nM 5-OP-RU in the presence of nanobodies or control anti-MR1 antibody 26.5 at 10 mg mL^-1^. right: Cumulative data from n=2 healthy donor PBMC samples as per plots in D, at varied concentrations of 5-OP-RU. Open circles are donor 1, Closed circles are donor 2. **E.** Bar graph showing CD69 expression on SKW-3.MAIT cells after co-culture with C1R.wt cells plus different MOI of DH5α *E. coli* or 5-OP-RU, with or without blocking reagents. Data depicts n=2 technical replicates and is representative of n=2 independent experiments. **F.** Flow cytometric histogram overlays showing staining of mouse M12 cells with P1G5 clones C11 and D9 after treatment with Ac-6-FP or 5-OP-RU. Median fluorescence intensities are depicted next to peaks. Data is presentative of n=2 independent experiments. **G.** Left: Representative flow cytometric pseudocolour plots gated on TCRβ^+^ cells after splenocytes were cultured for 6 days with 5-OP-RU in the presence of blocking reagents. Right: Cumulative data from n=5 mice across n=3 independent experiments. All error bars depict standard error of the mean.

Given the supraphysiological levels of MR1 in C1R.MR1 cells, it was next assessed whether nanobody blockade extended to C1R.wt cells expressing endogenous (low) levels of MR1. Here, both C11 and D9-Fc bound 5-OP-RU-pulsed C1R.wt cells (**Figure 4B**), and blocked SKW-3.MAIT TCR activation by 5-OP-RU-treated C1R.wt cells (**Figure 4C**). In these assays, clone C11 outperformed D9, blocking SKW-3.MAIT cell approximately 2.5x more potently at this matched nanobody concentration. To determine whether C11 could block primary human MAIT cell activity, peripheral blood mononuclear cells (PBMCs) were incubated *in vitro* with 5-OP-RU for 6 days, and the MAIT cell response determined by expansion and cell trace violet (CTV) dilution at the end of culture. Both 26.5 and C11-Fc fully blocked MAIT cell proliferation (**Figure 4D**). Whether this blocking activity extended to bacterial-derived ligands was then tested, whereby C1R.MR1 cells were incubated with fixed *E. coli* at graded multiplicities of infection (MOI) and co-cultured with SKW-3.MAIT cells. C11-Fc blocked SKW-3.MAIT cell activity across all tested MOIs, similar to 26.5 (**Figure 4E**).

The ability to use this nanobody in mouse models would provide a major benefit to testing the efficacy of C11-based therapeutics in preclinical mouse models. Accordingly, the crossreactivity of the nanobodies was tested with mouse cells. Both C11 and D9 bound 5-OP-RU-pulsed M12 mouse antigen-presenting cells, but not those treated with Ac-6-FP, as evidenced by the positive shift in MFI by 269 and 314 from baseline, respectively (**Figure 4F**). In mouse splenocyte cultures, 5-OP-RU treatment induced substantial MAIT cell proliferation across 6 days, reaching 7-20% of total TCR-b^+^ cells at the end of culture in presence of an isotype control or in absence of antibody (no block; **Figure 4G**). C11, D9, as well as 26.5 each completely abrogated this proliferation (**Figure 4G**). Together, these data demonstrate that C11 and D9 nanobodies block MAIT cell activation in an antigen-specific manner.

### In vivo activity of MR1-5-OP-RU-specific nanobody clone C11

To determine whether nanobody-mediated blockade could be effective *in vivo* applications, a systemic MAIT cell activation protocol was used whereby synthetic 5-OP-RU was administered intravenously. In this model, MAIT cells rapidly upregulate CD69 and CD25 in response to antigen, with activation detectable within two hours. Mice were injected intravenously with C11-Fc or clone 26.5 mAb 24 hours prior to 5-OP-RU administration, and organs were harvested two hours post-5-OP-RU injection for analysis (**Figure 5A**). At baseline, with no ligand injection, MAIT cells in the spleen and lung were largely negative or low for CD69 and CD25, whereas the majority of liver MAIT cells were constitutively CD69^+^ (**Figure 5B**). In response to 5-OP-RU treatment, strong upregulation of activation markers was observed across all tissues tested, where liver MAIT cells upregulated CD25 and those in the lung and spleen upregulated both CD25 and CD69 (**Figure 5B, C**). Pretreatment with clone 26.5 partially suppressed this activation in the spleen and lung, but had no effect in the liver. In contrast, C11-Fc fully blocked CD25 and CD69 upregulation in the liver and lung, and substantially reduced activation in the spleen (**Figure 5B, C**). These results demonstrate that bivalent C11-Fc can suppress MR1-5-OP-RU-driven MAIT cell responses *in vivo*.

**Fig. 5.**
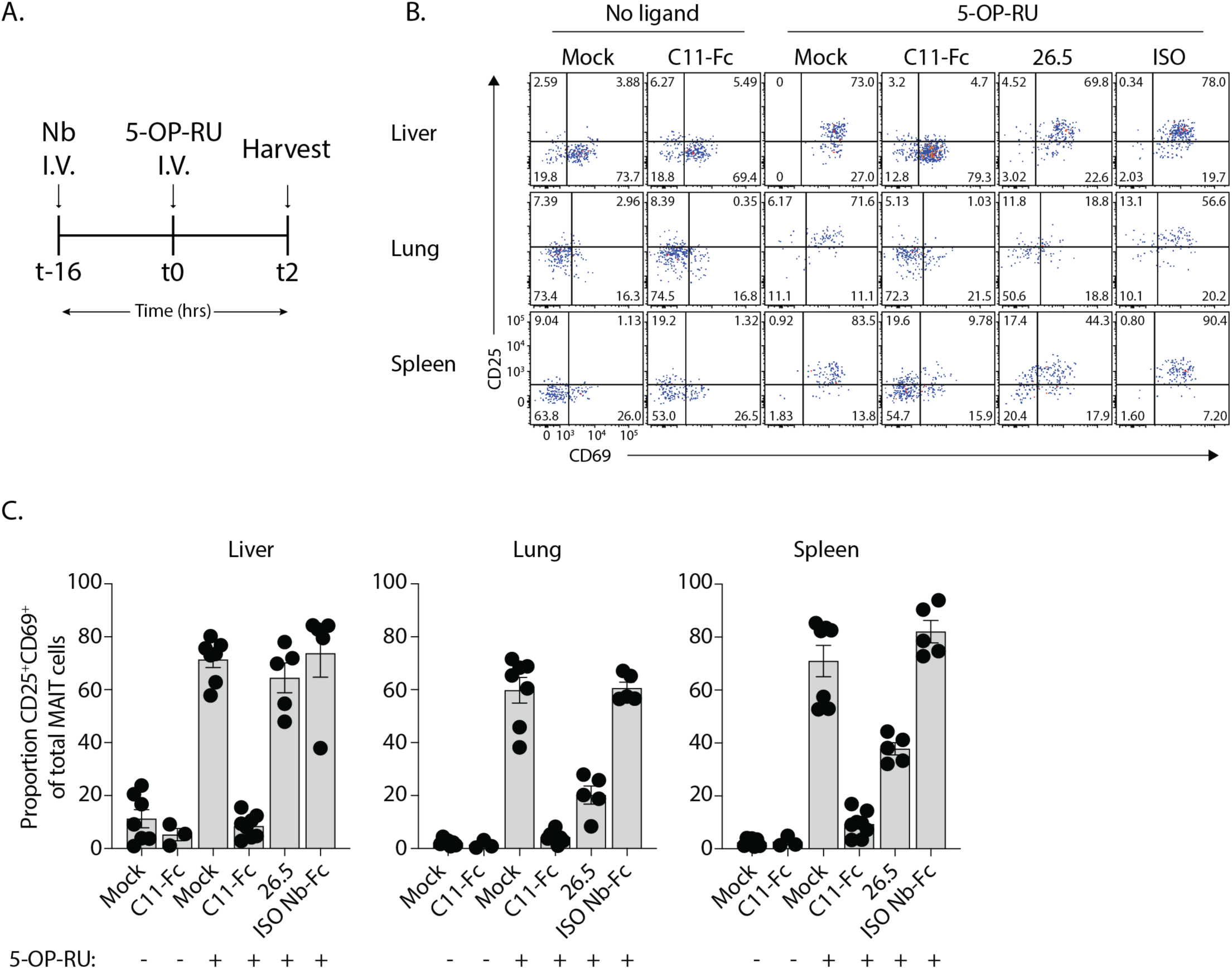
*In vivo* nanobody blockade of antigen-driven MAIT cell activation. **A.** Schematic of mouse challenge protocol. **B.** Representative flow cytometric pseudo-colour plots showing CD69 and CD25 expression on TCRβ^+^ MR1-5-OP-RU tetramer^+^ MAIT cells in liver, lungs and spleens from C57BL/6J mice 2 hours after IV injection of 5-OP-RU or PBS (Mock), with or without pretreatment with C11-Fc or 26.5 anti-MR1 antibody control. Numbers on plots represent percentages of quadrant gates. **C**. Bar graphs showing CD69, CD25 or co-expression on liver lung and spleen MAIT cells as per B from n=7 mice for PBS/PBS and PBS/5-OP-RU, n=8 mice for C11/5-OP-RU, n=5 mice for 26.5/5-OP-RU and ISO/5-OP-RU and n=3 mice for C11/PBS across n=3 independent experiments. Error bars depict standard error of the mean.

### Generation of bispecific antibodies targeting MR1-antigen complexes

Given the antigen-specificity and functional potency of the C11 nanobody, C11 may therefore have potential to specifically target cells furnishing 5-OP-RU in an immunotherapeutic context. The antigen-inducible upregulation of MR1 provides a potential opportunity to selectively target cells that actively present MR1-restricted ligands, including those infected with bacteria or tumour cells selectively presenting synthetic antigens like 5-OP-RU.

To test this concept, bispecific antibodies were generated, linking C11 to a widely characterised and clinically relevant clone of anti-human CD3, clone OKT3. These constructs were designed as single-chain diabodies (scDbs) incorporating the single-domain C11 nanobody in place of a conventional Fv region and joined via flexible linkers to the variable light and heavy chains of OKT3 in one contiguous polypeptide chain (**Figure 6A**). Two construct orientations were tested, one in which C11 was expressed at the C-terminal (OKT3xC11) and one at the N-terminal (C11xOKT3). Size-exclusion chromatography revealed two active peaks for each construct, both of which showed identical SDS PAGE gel migration patterns under both reducing and non-reducing conditions. The first peak corresponded to the expected size of monomers, whereas the second peak likely represented non-covalently-linked dimers (or multimer), consistent with mispairing of OKT3 heavy and light chains between two separate polypeptides (**Figure 6B**). All four proteins retained C11 binding specificity, validating an intact C11 domain, staining C1R cells pulsed with 5-OP-RU but not with Ac-6-FP or no ligand. The putative dimers had higher-intensity binding, in line with their higher valency (**Figure 6A-B**). Similarly, each protein efficiently labelled CD3 on TCR-transduced SKW-3 cells regardless of TCR identity, with the dimers providing brighter staining relative to the monomers (**Figure S5C-D**). These proteins also labelled T cells in human PBMC samples, with a similar staining hierarchy (**Figure S5E-G**).

**Fig. 6.**
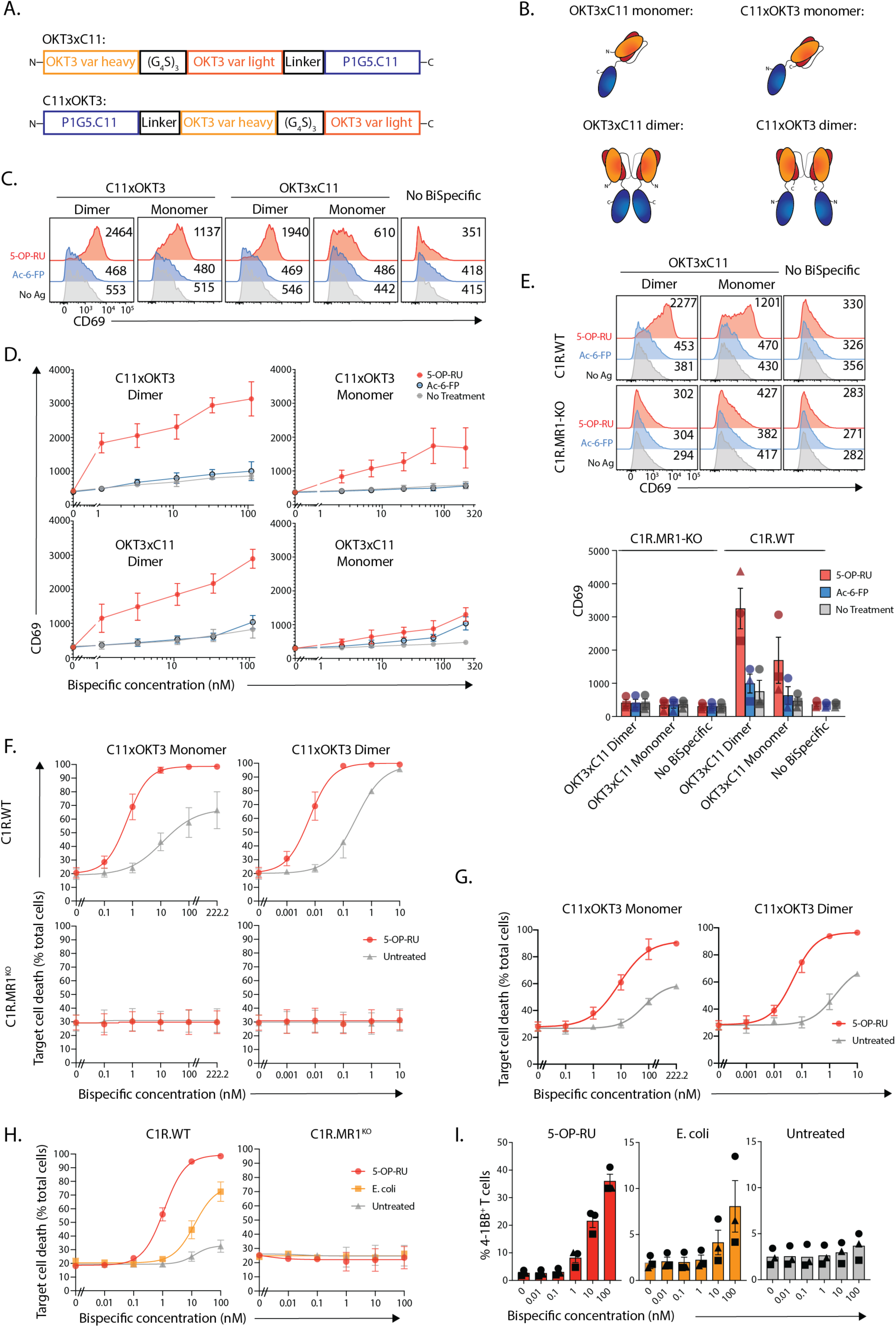
Development of bispecific antibodies targeting MR1-5-OP-RU complexes. **A.** Schematic of bispecific antibody constructs. **B.** Cartoon of Bispecific antibodies. **C.** Representative flow cytometric histogram overlays showing CD69 on SKW-3 cells expressing an MHC I-restricted TCR (SKW-3.TCR) after 18 hr co-culture with 5-OP-RU (10 mM; red), Ac-6-FP (500 mM; blue) or no antigen (grey)-pulsed wildtype C1R cells in the presence of bispecific antibodies at 1 µg mL^-1^. Numbers next to peaks depict median-fluorescence intensity (MFI) of CD69 staining. **D.** Top: Line graphs showing CD69 on SKW-3.TCR cells after co-culture as in C, in the presence of titrated concentrations of bispecific antibodies. **E.** Representative flow cytometric histogram overlays showing CD69 on SKW-3.TCR after co-culture as in C, using wildtype or MR1^-/-^ C1R cells. Numbers next to peaks depict median-fluorescence intensity of CD69 staining. **F-H.** Line graphs showing target cell death after co-culture of primary CD8+ T cells with **F.** C1R.wt or C1R.KO cells or **G**. SKBR3 cells in presence of 5-OP-RU, or **H.** C1R.wt or C1R.MR1KO cells with 5-OP-RU or DH5a *E. coli*, each with titrated concentrations of bispecific antibodies as labelled, or C11 OKT3 monomer in H. **I.** Bar Graphs showing % of T cells from co-cultures in H expressing 4-1BB after culture. Datapoints in D, F-H are mean of n=3 independent experiments, except C1R.MR1KO experiments represent a mean of n=2 independent experiments, whereas datapoints in E and I depict each of n=3 independent experiments. All error bars represent standard error of the mean.

To assess whether these bispecific antibodies were bioactive, C1R cells were pulsed with 5-OP-RU or Ac-6-FP, and then labelled with titrated concentrations of bispecific antibodies, before co-culturing them with SKW-3 cell lines expressing an MHC-I-restricted TCR (**Figure 6C-D**). Here, only those cells treated with 5-OP-RU elicited activation on the SKW3.TCR line by measure of CD69 expression. Activation was strongest with the C11xOKT3 dimer, followed by the OKT3xC11 dimer, with their monomeric forms substantially weaker and C11xOKT3 again outperforming OKT3xC11 (**Figure 6C-D**). In each instance, responses were 5-OP-RU specific with minimal activation in response to Ac-6-FP or no antigen, except at the higher concentrations of bispecific antibodies. This activation was strictly MR1-dependent, as MR1-deficient C1R cells failed to induce any response above baseline in the absence of bispecific antibodies (**Figure 6E**). Next, a similar assay was used to assess whether these bispecific antibodies could drive primary T cell-mediated cytotoxicity. C1R.wt cells (**Figure 6F**) and a breast tumour cell line, SKBR3 cells (**Figure 6G**), were pulsed with 5-OP-RU and labelled with titrating doses of C11xOKT3 monomer or dimer, then co-cultured with primary human CD8^+^ T cells. For both cell lines, both C11xOKT3 monomer and dimer induced potent 5-OP-RU-dependent killing, with the dimer consistently outperforming the monomer at lower concentrations (**Figure 6F**). Some antigen-independent killing occurred at high doses, especially with the dimer, but this was still MR1-dependent, as no killing was seen in MR1-deficient cells (**Figure 6F**). Finally, bispecific antibodies were tested for their ability to direct primary T cells to kill bacteria-infected cells. C1R.wt or C1R.MR1^KO^ cells were treated with *E. coli* or 5-OP-RU and used in similar assays (**Figure 6H**). Here, specific C1R cell death was observed in response to *E. coli* only in MR1-sufficient target C1R cells, and this correlated with reciprocal CD8^+^ T cell activation as measured by upregulation of early activation marker 4-1BB (**Figure 6I**). Thus, C11-based bispecific antibodies can redirect CD8^+^ T cells to kill antigen-treated tumour cells as well as those harbouring bacterial ligands, supporting the broader concept that MR1-antigen presentation can be exploited for selective immune redirection using nanobody-based bispecific antibodies.

## Discussion

This study demonstrates the potential for targeting MR1–antigen complexes using nanobodies, identifying clone C11 as a potent and specific binder to MR1–5-OP-RU. We show that yeast-display can generate nanobodies with fine specificity for MR1-antigen complexes, overcoming the challenge of recognising small, buried epitopes. The lead clone, C11, binds human and mouse MR1 in a 5-OP-RU–dependent manner and, following affinity maturation, achieves low nM affinity while preserving 5-OP-RU antigen specificity. Using a combination of crystallography with mutant MR1 protein analysis, we determined that mechanistically, C11 binds to residues flanking the A′ pocket, exploiting a minimal surface near the MAIT TCR footprint. This binding mode supports its capacity to selectively block MR1-dependent MAIT cell activation in both human and mouse systems. Notably, in contrast to clone 26.5, the nanobody had little impact on 5-OP-RU-independent MR1-reactive T cell responses, suggesting more targeted therapeutic potential than a pan-MR1 blocking approach.

To further demonstrate the immunotherapeutic potential of these nanobodies, we used one of the clones to generate bispecific antibodies capable of redirecting T cell responses toward cells displaying MR1-5-OP-RU complexes. This represents the first example of a TCRm bispecific antibody targeting a monomorphic antigen-presenting molecule, providing proof-of-principle for this approach. Compared to conventional TCRm platforms targeting polymorphic peptide–HLA complexes, MR1 possesses several intrinsic advantages, chief among them its limited allelic variation across individuals that enable universal therapeutic applicability. In addition, MR1 remains intracellular under steady-state conditions and only traffics to the cell surface upon antigen binding(*9*), thereby potentially limiting off-target effects. Furthermore, the ability to selectively target MR1 only with the 5-OP-RU antigen should also reduce the risk of inhibiting other MR1-reactive T cells that are 5-OP-RU independent and play important roles in the immune system(*22*).

There are multiple settings in which these tools could be deployed. Firstly, the riboflavin biosynthesis pathway, and hence production of 5-A-RU, is highly conserved across diverse microbial species, including clinically important pathogens(*7*). In infections where MAIT cell responses are insufficient or evaded, these bispecifics could compensate by recruiting a broad T cell response to eliminate infected cells. One potential application lies in latent *Mycobacterium tuberculosis*, where immune activation is limited but MR1 ligand production may persist. Alternatively, tumour cells could be targeted with a synthetic antigen bearing a ribityl-epitope to fine-tune and induce surface MR1-5-OP-RU expression, enabling a ‘bait-and-kill’ approach. In a similar manner, α-galactosylceramide (αGalCer) has been delivered directly into solid tumours to activate type I natural killer T (NKT) cells to elicit a localised anti-tumour response(*23*). We demonstrated that C11 is bioactive *in vivo*, as measured by blockade of MAIT cell activation by 5-OP-RU, and which provides promising preliminary data on the stability, tissue penetration and ability of C11 to bind MR1-antigen complexes *in vivo*. Future studies in mouse tumour or infection models will help determine the therapeutic potential of these nanobodies in disease settings. While the role of the tumour microbiome is an ongoing area of research(*24, 25*), microbial-based immunotherapies have shown pre-clinical promise in modulating the tumour microenvironment, including delivery of nanobody-secreting strains of *E. coli* to tumours(*26, 27*). Similar strategies could potentially be adapted using 5-A-RU sufficient bacteria engineered to secrete C11-based TCRm bispecific antibodies, thereby inducing local MR1-antigen expression within tumours.

Beyond bispecific antibodies, C11 could also be engineered for other therapeutic applications. For instance, C11 nanobody blockade of MR1 could be used to suppress pathological antigen-driven MAIT cell activation in inflammatory or infectious settings, as previously shown in a mouse model of *Helicobacter pylori* where monoclonal antibodies could suppress MR1-antigen-driven pathology(*28*). The specific targeting of MR1 only when bound-to microbially derived 5-OP-RU, as achieved with the nanobodies described here, would also minimise off target effects against other cells bearing MR1, and other MR1-restricted T cells that may play important immunological roles(*22*). Due to their small size and lack of an Fc domain, nanobodies have superior tissue penetration, conformational stability and short half-lives; pharmacological properties which may be advantageous for contexts requiring transient immunomodulation. C11 could also be adapted for payload delivery, for example, in antibody–drug conjugates targeting bacteria-infected cells(*15*), or engineered with an Fc domain, as was performed here, to extend half-life and *in vivo* persistence.

It is important to consider that 5-A-RU is produced not only by pathogens but also by commensal microbes, and conditions under which MR1 presents other antigens derived from 5-A-RU remain incompletely defined. Future studies investigating when and where MR1-5-OP-RU complexes arise, particularly under homeostatic or inflammatory conditions, will be helpful to gauge the risks of off-target effects in therapeutic contexts. In this regard, C11 represents a valuable tool where its cross-reactivity between humans and mice make it well-suited in tracing MR1-ligand complexes and determining the safety of such an approach in both murine models and human tissues.

This work establishes a conceptual and technical foundation for targeting MR1–antigen complexes with high precision. The yeast-platform applied here is readily adaptable, and opens the door to the rapid discovery of nanobodies binding emerging MR1 ligands, for example, the recently described carbonyl nucleobase adducts that are MR1-presented antigens for a population of non-MAIT T cells(*29*). These antigens are associated with cellular stress including in cancer cells, and therefore represent high-potential targets for biologics like the highly specific nanobodies described here. This approach could also be extended to other monomorphic antigen-presenting molecules. For example, HLA-E, HLA-F and HLA-G exhibit limited polymorphism and present conserved peptide antigens for T cell recognition, and CD1 family members that bind and present lipid antigens(*1*). Although nanobodies targeting CD1d have been described, their mechanisms of action are distinct to C11. Some bind CD1d and co-engage the NKT TCR to drive NKT cell responses^30^, while others bind in an antigen-independent manner and modulate CD1d antigen-presentation(*30, 31*). Antibodies and nanobodies binding CD1d-α-GalCer complexes have also been developed, though their therapeutic potential has not been explored(*32*). Similarly, a CD1c-restricted TCR that binds CD1c molecules presenting self-lipids in an antigen-independent manner has been affinity-enhanced and engineered as a bispecific engager, and demonstrated to redirect T cell responses toward CD1c^+^ leukemias(*33*). The platform established here could accelerate discovery of antigen-specific binders across these distinct monomorphic systems. There is great interest in generating peptide-MHC-specific binders, and a number of platforms have been established to fast-track their development, including the *de novo* generation of MHC-binding peptide scaffolds and pipelines employing artificial intelligence (AI) to generate display libraries of high potential binders to great effect(*34–38*). We show here that a naïve synthetic yeast display library can also be used to rapidly generate binders, without the need for AI-guided design and modelling steps, nor an alternative peptide platform. Nonetheless, C11 may serve as useful scaffold for directed evolution of related specificities, forming new mutant libraries to enhance the starting frequencies of clones with predilection for MR1-binding but varying antigen-specificity.

Collectively, this work establishes a flexible platform for the discovery of MR1-ligand binders, defines the first MR1–5-OP-RU–specific nanobody, and demonstrates its utility across biochemical, functional, and translational settings. More broadly, this work expands the TCRm antibody paradigm to monomorphic antigen-presenting molecules, introducing a new class of molecules with therapeutic potential and investigative reagents for the study and manipulation of the MR1-MAIT cell axis.

### Limitations of the study

This work serves as proof-of-concept for targeting MR1-antigen complexes using TCRm nanobodies, in this case, using clone C11 to target MR1-5-OP-RU complex as a model. Moreover, the utility of the nanobody is exhibited by demonstrating its ability to inhibit antigen-driven MAIT cell responses, as well as to redirect broad T cell responses toward tumours or infected cells. Further *in vivo* characterisation would be required to guage the efficacy and safety profile of such an approach in diverse models. For example, the possibility that microbiome-derived 5-OP-RU could be presented by MR1 at barrier tissues highlights the need for careful evaluation of on-target effects in physiological settings. These considerations are beyond the scope of this first study but will be important for future translation.

## Materials and Methods

### Study Design

The primary objective of this study was to develop nanobodies that bind MR1-5-OP-RU complexes in an antigen-dependent manner. Subsequent controlled laboratory experiments were designed to characterise the nanobody specificity, structural basis of binding, and functional utility, including bispecific engager. Randomisation and blinding were not relevant to this study.

### Human PBMC samples

Human peripheral blood mononuclear cells were isolated from buffy coats isolated from healthy blood donors in Melbourne, Australia and supplied by the Australian Red Cross after informed concent (agreement numbers: 17-08-VIC-16, 18-08-VIC-12, 20-10VIC-14, 22-11VIC-04 and 24-10VIC-15) in accordance with human ethics approval from the University of Melbourne Human Ethics Committee (ethics number: 30058).

### Mice

C57BL/6J colonies of either wild-type or *Cd1d^−/−^Tcrd^−/−^*(*39*) background were utilised. Mice were housed in the Department of Microbiology and Immunology Biological Resource Facility, University of Melbourne at the Peter Doherty Institute for Infection and Immunity. Age- and sex-matched controls were used where appropriate. All procedures conducted were in accordance with approved ethics from the University of Melbourne Animal Ethics Committee (ethics number: 24324), in line with the NIH Guide for the Care and Use of Laboratory Animals.

### Cell lines

Cells were cultured in complete media using either Dulbecco’s Modified Eagle Medium (DMEM) or Roswell Park Memorial Institute-1640 (RPMI-1640). The media were supplemented with 2% foetal bovine serum (FBS) and serum complement consisting of L-Glutamine (2 mM), non-essential amino acids (0.1 mM), HEPES (15 mM), penicillin/streptamycin (100 units mL^-1^ each), sodium pyruvate (1 mM), 2-mercaptoethanol (50 mM)). Cells were cultured at 37° C, 5% CO_2_ and were passaged every two to three days as required. Adherent cell lines underwent trypsinisation prior to passaging or harvest for cellular assays. C1R and M12 cells were obtained from JM Laboratory (University of Melbourne). SKBR3 cells were obtained from Associate Professor Paul Beavis (Peter MacCallum Cancer Centre). Jurkat-76(*40*) and SKW-3 reporter cell lines were previously described for M33-64(*19*), MBV28(*19*), MC.7.G.5(*41*), DGB129(*41*), G115(*42*), NKT15(*43*), TCR21(*44*) and OTN5(*45*), as well as C1R.CD1c(*46*), C1R.MR1(*19*), and C1R.MR1^KO^ (*41*).

### Isolation of murine lymphocytes

Spleen were harvested from mice and mechanically digested through a 30 µM nylon sieve into ice cold FACs buffer (2% FBS in PBS), and subjected to red blood cell lysis for 5 minutes at room temperature (RT) (Sigma-Aldrich). Liver and lungs were perfused with 10 ml ice-cold PBS immediately after euthanasia. Livers were mechanically digested through a 70 µM nylon sieve and underwent a density gradient spin with 33% Percoll solution (Cytiva). Lungs were harvested and minced in type III collagenase buffer (3 mg/ml; Worthington Biochemical Corporation in RMPI-1640 supplemented with 2% FCS), followed by incubation in a shaker incubator for 1 hour at 37°C. The liver and lung samples were then filtered through a 30 µM nylon sieve and underwent red blood cell lysis. Samples were finally resuspended in ice cold FACs buffer.

### Mouse ex vivo activation and proliferation assays

Splenocytes from either C57BL/6J wild type or *Cd1d^−/−^Tcrd^−/−^*mice(*39*) were processed as described above. The resulting suspensions were pooled for each respective strain and underwent a further histopaque-1083 (Sigma Aldrich) gradient spin to isolate viable cells prior to as per manufacturer’s instruction (Invitrogen) labelling, and then cultured in complete DMEM in the presence of rhu-IL2 (100U). Splenocytes were cultured for 4 days with 5-OP-RU (Provided by JYWM and DPF, University of Queensland). Cultures were harvested on either day 1 or day 4 post culture to assess early MAIT cell activation or proliferation respectively. Mouse MAIT cells were identified as B220^-^ TCRβ^+^, MR1-5-OP-RU tetramer^+^ cells. Activation was assessed via CD69 and CD25 status and proliferation was assessed via MAIT cell expansion and CTV^-/low^ cell frequencies.

### In vivo MAIT cell activation blockade

C57BL/6J wildtype mice were injected with Fc silenced C11-Fc D265A or anti-MR1 followed by synthetic 5-OP-RU after 16 hours. Mice were culled 2 hourspost 5-OP-RU administration. Lungs, spleen and livers were harvested, processed as described above and analysed via flow cytometry.

### Flow cytometry

Unless stated otherwise, cells were stained in antibody cocktails in FACS buffer consisting of phosphate buffered saline (PBS) supplemented with 2% (v/v) FBS for 20 minutes at RT, in the dark, washed once, fixed in PBS supplemented with 2% paraformaldehyde (PFA), washed once more and then acquired using a BD LSR Fortessa. Data was analysed using Flowjo Software (BD; Version 10). Antibodies clones, titres and vendors are included in **Supplementary Table S1**.

### Antigen synthesis

Synthetic 5-OP-RU was prepared as a solution in DMSO-d_6_, as previously described(*47, 48*). Its concentration was determined via quantitative NMR spectroscopy(*49*). This water-reactive antigen was diluted in aqueous buffer immediately before use.

### MR1 upregulation assays

Antigen-presenting cells were pulsed with 5-OP-RU (10 µM) or Ac-6-FP (500 µM; Schircks Laboratories) for 3 hours. Cells were then harvested and labelled with Nb-Fc proteins and controls, followed by staining with secondary goat anti-mouse IgG (PE) antibody. Staining was assessed via median fluorescence intensity (MFI) of PE via flow cytometry. Cells of interest were gated as single, viable cells, including GFP^HI^ for those lines expressing MR1.

### T cell reporter activation assays

SKW-3.TCR or Jurkat-76.TCR cells were co-cultured with CTV-labelled antigen-presenting cells in the presence of 5-OP-RU or infected with PFA-fixed DH5α *E. coli*, as well as blocking antibodies (10 mg mL^-1^ unless stated otherwise in figures). Cells were incubated for approximately 18 hoursbefore staining with LIVE/DEAD Fixable Near-IR Dead Cell Stain and anti-human CD69 before being acquired by flow cytometry. T cell reporters were identified by gating on viable GFP^HI^ cells.

### Bispecific antibody activation assays

For C1R co-culture assays, C1R cells were labelled with CTV as per manufacturer’s instructions (Invitrogen), and pulsed with 5-OP-RU (10 µM) or Ac-6-FP (500 µM) for three hours, selecting these doses for their comparable induction of MR1-surface upregulation. Cells were then washed twice and incubated with bispecific antibodies for 15 minutes at 4°C and then washed five times with complete media. Subsequently, cells were co-cultured for approximately 18 hours with a SKW-3.TCR line expressing an MHC I-restricted TCR, clone OTN5(*45*) at a ratio of 1:1. Cells were then harvested, stained with anti-human anti-CD69 BUV737, and a LIVE/DEAD Fixable Near-IR Dead Cell Stain and acquired by flow cytometry. TCR^+^ cells of interest were identified by gating on viable CTV^-^, GFP^HI^ cells. For plate-bound activation assays, bispecific antibodies were immobilised on a 96 well plate by incubation for at 37°C for 2 hours. Wells were washed with PBS and 100,000 T cell reporter cells were plated per well. Cells were incubated for approximately 18 hours before harvesting and staining as above, followed by acquisition by flow cytometry.

### Killing assays

Target cells were labelled with CTV and subsequently pulsed for 3 hours as above. For infection, C1R cells were CTV-labelled as per manufacturer’s instruction (Invitrogen) and incubated overnight with either 5-OP-RU (10 µM) or PFA-fixed DH5α *E. coli* at 100 MOI. After pulsing, cells were subsequently washed five times, incubated with bispecific antibodies, washed again, and co-cultured for approximately 18 hours with primary *in vitro* expanded CD8^+^ T cells at a 2:1 effector: target ratio. SKBR3 cells were harvested and stained with (7-AAD), anti-human CD3, and anti-human CD8a, whereas C1R cells were stained with (LIVE/DEAD Fixable Near-IR Dead Cell Stain), anti-human CD3, and anti-human CD8. Cells were then acquired by flow cytometry. Target cells were electronically gated on CTV^+^, CD3^-^, CD8^-^ cells whilst effector cells electronically gated on viable CTV^-^, CD3^+^, CD8^+^ cells.

### In vitro expanded CD8 T cells

PBMC were stained with anti-CD3 BUV395, anti-CD4 BUV496, anti-CD8 BUV805, anti-CD14 BV570, anti-CD19 APC-Cy7, anti-Va7.2 BV711, anti-gdTCR PeCy7, huMR1-5-OP-RU tetramer-BV421 and 7-AAD and subsequently FACS-sorted for 7AAD^-^, CD14^-^, CD19^-^, CD3^+^, CD8a^+^, CD4^-^, huMR1-5-OP-RU tetramer^-^T cells. Cells were then cultured in complete T cell media (as per complete media above, using a 1:1 mix of AIM-V and RPMI-1640 as a base rather than 100% RPMI-1640) with plate-bound anti-CD3 (clone OKT3) and anti-CD28 (clone 9.3) mAb in the presence of rhu-IL-2 (200 U mL^-1^, Peprotech). Stimulation was removed 48 hourspost-culture and T cells were cultured with rhuIL-2 supplemented media for 14 to 18 days. Purity (>95% CD8^+^ T cells) was confirmed via staining with relevant antibodies and flow cytometry analysis prior to cryopreservation. Prior to use, T cells were thawed and rested overnight in T cell media.

### Nanobody selection

The naive nanobody yeast display library was provided by Professor Andrew Kruse, Harvard Medical School, and previously described in detail(*14*). The selection strategy was similar to that we previously published(*50*). In brief, nanobody expression was induced in the naive library by replacement of glucose with galactose-supplementation in tryptophan knockout media (-Trp) for 48 hours. The library was first magnetically depleted by 30 minutes incubation with 100 nM human unloaded CD1c-(CD1c-endogenous; CD1c-endo) PE-Cy7 Tetramers and passing over a MACS LD column (Miltenyi Biotec) followed immediately by magnetic enrichment including 30 minutes incubation with 100 nM huMR1-5-OP-RU PE tetramers and passing over a MACS LS column (Miltenyi Biotec). Enriched cells were recovered in glucose-supplemented -Trp media for 24 hours before being transferred into galactose-supplemented - Trp media for 48-72 hours prior to a subsequent depletion/enrichment round. Three rounds of magnetic depletion/enrichment were performed before one round with flow cytometry-based enrichment. Here, cells were first labelled for 30 minutes at 4°C with 100 nM human CD1c-endo PE-Cy7 tetramers, followed by avidin and biotin blocking (Dako) for 30 minutes each at 4°C as per manufacturer’s instructions, before incubation with staining cocktail including LiveDead Near InfraRed (1:500), anti-HA AF647 (1:200) and 100 nM huMR1-5-OP-RU PE tetramers for 30 minutes at 4°C. Yeast cells were then FACS-sort enriched using a MoFlo Astrios (Beckman Coulter) for viable CD1c-endo tetramer^-^, MR1 tetramer^+^ yeast cells. Cells were sorted directly into -Trp media supplemented with glucose and recovered for 72 hours. For single cell colony sorts, galactose-induced cells were first labelled with 100 nM human CD1c-endo PE-Cy7 tetramer and huMR1-6-FP PE-Cy7 tetramers for 30 minutes at 4°C, followed by avidin and biotin blocking as above before a second labelling step with staining cocktail including LiveDead Near InfraRed (1:500; Invitrogen), anti-HA AF647 (1:50, Biolegend) and 100 nM huMR1-5-OP-RU PE tetramers for 30 minutes at 4°C. Nanobody induction was measured prior to enrichments by staining with Live-Dead Near Infrared (1:500) and anti-HA AF647 (1:200) followed by flow cytometric analysis. Enrichment-efficiency and specificity was measured between enrichment rounds after 48 hoursgalactose induction by staining with Live-Dead Near Infrared (1:500), anti-HA AF647 (1:200) and 100 nM CD1c-PE-Cy7 or huMR1-5-OP-RU PE tetramers or streptavidin alone. Cells were then single cell FACS-sorted into 96-well plates with 100 ml glucose-supplemented -Trp media using a MoFlo Astrios for viable nanobody (HA)-expressing yeast that were PE^+^PECy7^-^ tetramer single positive. After 72 hours recovery and colony growth, nanobody expression was induced on yeast colonies in 96-well plates with 100 ml galactose -Trp media for 48 hours. Cells were then stained for 30 minutes with Live-Dead Near Infrared (1:500) and anti-HA AF647 (1:200) followed by staining for 30 minutes with 100 nM huMR1-5-OP-RU PE tetramers, huMR1-6-FP PE tetramers, mouse MR1-5-OP-RU PE tetramers, mouse MR1-Ac-6-FP PE tetramers or streptavidin-PE alone. Plates were then acquired with a LSR Fortessa X-20 (BD) or CytoFLEX. Colonies were cryopreserved by resuspending yeast pellets in glucose-supplemented -Trp media with 15% glycerol in 96-well plates and stored at -80°C.

### Affinity maturation library generation

Affinity maturation was performed as previously described(*51*). Here, pYDS plasmid encoding P1G5 or P2A3 nanobody clones were isolated from yeast colonies via yeast miniprep (Zymo) and plasmids transformed into DH5a E. coli via electroporation before bacterial miniprep (Zymo) and sequence verification. Error prone PCR was then performed on plasmids using AB_MFAL_fwd (GTTCAATTGGACAAGAGAGAAGCT) and AB_HA_rev (GAACATCGTATGGGTAGGATCC) according to manufacturer’s instructions (Agilent GeneMorph II Random Mutagenesis kit) using 40ng, 15ng, 5ng or 2ng template DNA with 2 minutes at 95°C followed by 30 cycles of 30 sec at 95°C, 30 sec at 50°C and 1 minutes at 72°C followed by a final 10 minutes at 72°C. PCR products were then cleaned up using a Promega Wizard SV gel and PCR cleanup kit. Each PCR product was then amplified across 12x 50 ul reactions per PCR product library using Kapa HIFI Hotstart ReadyMix (Roche) with AB_Lib_fwd (ATCGCTGCTAAGGAAGAAGGTGTTCAATTGGACAAGAGAGAAGCT) and AB_Lib_rev (GGGTGAGGATGTTTGAGCGTAATCTGGAACATCGTATGGGTAGGATCC) primers as per manufacturer’s instructions with 3 minutes at 95°C followed by 30 cycles of 20 sec at 98°C, 15 sec at 65°C and 15 sec at 72°C, and a final 1 minute at 72°C. PCR products were separated by gel electrophoresis, appropriate bands excised and DNA recovered using DNA gel recovery kit (Zymo). *Sacchromyces cerevsiae* BJ5465 yeast (In Vitro Technologies; ATC208289) were electroporated with 15 mg pYDS plasmid digested with NheI and BamHI with 75 mg nanobody mutant library DNA consisting of 1:1:1:1 ratio of the 4x libraries generated above. Electroporations were distributed across 9 cuvettes (Cell Projects, #EP-202), and performed using a GenePulser Xcell (BioRad) using the square wave protocol at 500V for 15 ms. Yeast were recovered for 45 minutes in 20 ml YPD media at 30°C with no shaking. Aliquots were plated on Agar plates for cell counts to estimate library size (P1G5 = 108 million; P2A3 = 113 million). Remaining yeast were transferred to glucose-supplemented -trp media, cultured for 72 hoursat 30°C, 220 rpm before cryopreservation.

### Affinity maturation yeast selections

For each of the P1G5 and P2A3 mutant libraries, 5e8 Nb-induced yeast were stained with 190 nM biotinylated huMR1-5-OP-RU monomer for 30 minutes at RT. After three washes, yeast were then stained with cocktail containing Streptavidin-PE (Molecular Probes) at 1 mg ml^-1^, anti-HA AF488 (1:200) and Live-Dead Near Infrared (1:500) for 15 minutes at RT. Cells were then washed twice prior to FACS-sorting for HA^+^ huMR1-5-OP-RU^+^ viable yeast using a MoFlo Astrios (). Cells were recovered in glucose-supplemented -Trp media for 24 hr before 72 hr induction in galactose-supplemented -Trp media prior to the next selection round. Two further rounds of enrichment were performed as above, reducing the concentration of huMR1-5-OP-RU to 19 nM and 1.9 nM in subsequent rounds. Cells were then sorted into single cell colonies as per the naïve library sorts, and stained as above using 1.9 nM huMR1-5-OP-RU. 44 colonies were screened for huMR1-5-OP-RU, huMR1-Ac-6-FP and mouse MR1-5-OP-RU monomer staining at 1.9 nM as above, cryopreserved and sequenced.

### Yeast colony nanobody sequencing

For non-affinity matured yeast colonies, selected colonies were inoculated in 5 ml glucose-supplemented media at 200 rpm, 30°C for 48 hours. Yeast was then streaked onto glucose-supplemented -Trp Agar plates. After 48 hours, colonies were picked and miniprepped (zymo yeast miniprep kit II). Nanobody-encoding DNA was then PCR-amplified from miniprepped plasmids. PCR products were separated via gel electrophoresis and DNA bands corresponding to amplified DNA excised from the gel and DNA purified using a DNA gel purification kit (Zymo Research). DNA was digested with NheI and BamHI and fragment ligated into pHLSec plasmid encoding a Nb-Fc fusion. pHLSec plasmids were then sequenced via sanger sequencing (AGRF) and midiprepped or maxiprepped (Zymo Research) for protein expression.

For affinity matured yeast colonies, selected colonies were cultured in 200 ml glucose-supplemented -trp media in wells of a 97-well plate before 5 ml of each colony was transferred to wells of a PCR plate. Yeast were subsequently digested 3 hours at 37°C after addition of 100 ml of digestion mix containing 10% sucrose (w/v), 50 mM Tris 8.0, 10 mM EDTA, 1:1000 2-ME and 0.2 mg/ml Zymolyase 20T before inactivation for 5 minutes at 98°C in a thermocycler (Eppendorf). Samples were then diluted 1:10 in nuclease-free water and 2 ml used as template in a PCR reaction using pYDS649HM_fwd (GTTTTGTTCGCTGCTTCTTCTGC) and pYDS649HM_rev (GGTAGATTTACTAGGCGATGAGG) primers with 3 minutes at 95°C followed by 35 cycles of 20 sec at 98°C, 30 sec at 65°C and 1 minutes at 72°C with a final 1 minutes at 72°C. 5 ml PCR product was treated with 2 ml ExoSAP-IT (Thermo Fisher Scientific) for 30 minutes at 37°C followed by 15 minutes at 80°C and sequenced by Sanger sequencing (Australian Genome Research Facility; AGRF) using the pYDS649HM_fwd primer.

### Protein expression, purification and labelling

Biotinylated huMR1-5-OP-RU, huMR1-6-FP, mouse MR1-5-OP-RU and mouse MR1-Ac-6-FP tetramers were produced in house as previously described(*7*), using a 6xHIS-tagged construct with AVI-tag(*52*). Here, MR1-Antigen complexes were refolded from inclusion-body preparations by oxidative refolding between MR1 heavy chain, b2-microglobulin and ligands followed by dialysis, Ni-NTA purification, enzymatic biotinylation and size-exclusion chromatography gel filtration. Biotinylated human CD1c-endo tetramers were produced in house as previously described(*46*) by transient expression in mammalian Expi293F cells via expifectamine as per manufacturer’s instructions (Thermo Fisher Scientific), followed by purification as per MR1 proteins. Biotinylated MR1 and CD1c were tetramerised by sequential equimolar addition of streptavidin-PE or streptavidin-PE-Cy7 (BD Biosciences) at a 4:1 molar ratio of monomer:streptavidin.

For MR1 proteins used in crystallography and SPR, soluble human and mouse wild-type MR1-5-OP-RU and MR1-Ac-6-FP proteins were generated in house, as described previously(*7, 8*). In brief, MR1-β2m monomers were folded from *E. coli* inclusion bodies in the presence of MR1 ligands via oxidative refolding, dialysed and purified using sequential anion exchange (DEAE Sepharose, GE Healthcare), size-exclusion (Superdex 75pg 16/600 gel filtration column; Cytiva) and strong anion-exchange (MonoQ, GE Healthcare) chromatography. MR1-antigen proteins were analysed for purity by SDS-PAGE.

Monomeric nanobodies were produced via bacterial periplasmic expression as previously described(*14*). First, codon-optimised dsDNA fragments encoding nanobody protein with an N-terminal PelB sequence and C-terminal 6xHIS-tag (IDT) were cloned into pET-30 plasmid. Then, plasmids were transformed into BL21 E. coli and colonies were grown overnight at 37°C, 180 rpm in 10 ml starter cultures of Terrific Broth supplemented with 30 mg ml^-1^ kanamycin and 24 mml^-1^ chloramphenicol. Cultures were then used to inoculate 1 L flasks of pre-warmed kanamycin and chloramphenicol-supplemented Terrific Broth. After approximately 4 hours at 37°C, 180 rpm, when the OD reached 0.6-0.8, protein expression was induced by addition of 1 mM IPTG (BioVectra). Following induction, flasks were shaken at 25°C at 180 rpm for approximately 16 hours. Bacteria were then pelleted and resuspended in 100 mL 0.5 M sucrose, 0.2 M Tris, 0.5 mM EDTA at pH 8.0, 4°C. Bacteria were then osmotically shocked by addition of 200 mL H_2_0 with 45 minutes stirring at 4°C. Lysate was then adjusted to 150 mM NaCl, 2 mM MgCl_2_ and 20 mM Imidazole before being pelleted at 20,000g for 20 minutes at 4°C. Supernatant was then harvested, 0.2 mm filtered and purified by Ni-NTA followed by size-exclusion chromatography gel filtration using a Superdex 75pg 16/600 gel filtration column (Cytiva). Nanobodies were 0.2 mm sterile filtered and stored in PBS at –80°C until further use.

Fc-linked nanobodies fused full-length nanobodies at the C-terminus to mouse IgG1 from the hinge region to the end of the CH3 domain via a GS-linker to generate Nb-Fc fusion proteins where the linking section is encoded by …TVSS-GSGSG-CKPCIC. Where described, some constructs included a D265A mutation in the CH3 domain to reduce Fc-binding. Isotype control protein nanobody clone 5E7(*53*) was also produced in house. DNA encoding the nanobody was cut between NheI and BamHI directly from the DNA amplified from yeast colonies, and ligated into an existing pHLSec mammalian expression plasmid encoding a Nb-Fc template. scDb constructs was designed as previously described, with two orientations, either N-terminal OKT3 scFv with heavy and light chain separated by (G_4_S)_3_ linker, followed by a GTGAS linker prior to the nanobody sequence and C-terminal 6xHis-tag , or N-terminal nanobody followed by GSGTG linker and C-terminal OKT3 scFv and 6xHis-tag . Both Nb-Fc and scDb proteins were expressed by transient transfection of Expi293F cells using expifectamine. Cells were cultured for 6 days as per manufacturer’s instructions before cells were pelleted and supernatants were harvested and 0.2 mm filtered ready for purification. Nb-Fc proteins were purified by protein-A HiTrap Prism A MabSelect columns (Cytiva) followed by size-exclusion chromatography gel filtration using a Superdex 200pg 16/600 gel filtration column (Cytiva). scDbs were purified by Ni-NTA HisTrap FF columns (Cytivia) followed by size-exclusion chromatography gel filtration using a Superdex 75pg 16/600 gel filtration column (Cytiva). Both proteins were then sterile 0.2 mM filtered and stored in PBS at –80°C prior to use.

### Surface plasmon resonance (SPR)

Reagents for amine coupling: (1-ethyl-3-(3-dimethylaminopropyl) carbodiimide hydrochloride (EDC), *N*-hydroxysuccinimide (NHS)) and ethanolamine) as well as the Series S CM5 sensor chip were acquired from Biacore, GE Healthcare.

SPR binding experiments were performed on a Biacore T200 instrument at 25°C, in working buffer, 10 mM HEPES pH7.5, 150 mM NaCl 0.005% surfactant P20 (HBS-150). To control the background signals, no protein was immobilised on the designated reference flow cell. The antibody or nanobody of interest was coupled to the CM5 sensor chip (Biacore) by standard amine coupling, using the standard protocol(*54, 55*). Following injection of EDC/NHS across all flow cells, immobilisation of each ligand to the sensor chip was achieved using 10 mM sodium acetate pH 4.7. The surface density of immobilised ligand was kept consistent per CM5 chip, with wild-type nanobody (∼2000 resonance units; RU), C11, and 26.5 mAb each immobilised at ∼300-400 RU, respectively. The analytes (human and mouse MR1-5-OP-RU, hMR1-Ac-6-FP) were serially diluted in HBS-150 (0-40 µM) and passed over all flow cells at 30 µL min^-1^, with measurements taken in duplicate (n=2). Following each run, the sensor surface was regenerated with gentle regeneration buffer (Thermo Fisher Scientific), dissociation would not otherwise reach baseline. The SPR sensorgrams, equilibrium curves, and steady state affinity K_D_ values (µM) were prepared in T200 evaluation software and using GraphPad Prism.

### Crystallisation, structure determination and refinement

Soluble C11 nanobody was mixed with MR1-5-OP-RU or MR1-Ac-6-FP proteins at 1:1 ratio in the buffer (10 mM Tris-HCl, 150 mM NaCl pH 8.0) and the tertiary complexes were purified by size exclusion chromatography. C11-MR1-Ags complexes was concentrated to final concentration of 10mg/ml. To identify suitable crystallisation conditions, sparse matrix screening was performed involving the commercially available screens PACT Premier, JCSG+, ProtComplex, Morpheus, MorpheusII, Wizard classical 1&2, JBScreen Classic HTS I and JBScreen Classic HTS II. For this, protein complexes (10, mg/mL) were mixed with reservoir solution in a 1:1 volume ratio (200 nL:200 nL) and subjected to hanging-drop vapour diffusion at 20°C. Initial crystals of C11-MR1-5-OP-RU and C11-MR1-Ac-6-FP appeared after 2-6 days with a precipitant consisting of 100 mM HEPES 7.3 pH, and 4.2 M NaCl. After manual grid optimization around this original condition, single plate shaped crystals were grown over a week against a reservoir solution of 100 mM HEPES 7.5 pH, and 4 M NaCl at 20°C.

### X-ray diffraction data collection and structure determination

C11-MR1-5-OP-RU and C11-MR1-Ac-6-FP crystals were flash-frozen in liquid nitrogen after quick washing in reservoir solution. X-ray diffraction data were collected at 100 K on the Australian Synchrotron at MX2 beamline(*56*). Diffraction images were indexed, integrated and scaled using XDS(*57*) and further processed and analysed using programs from the CCP4 suite(*58*) and the Phenix software package(*59*). The C11-MR1-5-OP-RU and C11-MR1-Ac-6-FP structures were determined by Molecular Replacement using PHASE(*60*), where modified nanobody (PDB ID; 4S11) and MR1-_2_m (PDB ID; 4L4T) were used as search models. Afterwards, an initial run of rigid body refinement was performed with Phenix.refine(*59*) and the CDRH loops of C11 were subsequently rebuilt using the program COOT(*61*). Iterative rounds of model building using COOT and refinement using Phenix.refine were performed to improve the model. The Grade Webserver and Phenix tools were used to build and to generate ligand restraints(*58*). The structure was validated using MolProbity(*62*) and graphical representations were generated using PyMOL Molecular Graphics System, Version 2.2 (Schrödinger, LLC, New York, NY). The quality of the structure was confirmed using the Research Collaboratory for Structural Bioinformatics Protein Data Bank Data Validation and Deposition Services. The total interface area was evaluated by PISA analysis(*63*) and the contacts were analysed by the Contact program, both form the CCP4 suite. Statistics on the data collection and the final model are summarized in **Supplementary Table S2**.

### Statistical Analysis

Statistics including the description of error bars and experimental and technical replicates are described in detail in the relevant figure legends.

## Acknowledgements

Funding: HFK was supported by an ARC Discovery Early Career Research Award (DE220100830) and acknowledges grant support from an NHMRC Ideas Grant (2030162). NAG was supported by an ARC Discovery Early Career Research Award (DE21010070) and now an NHMRC Emerging Leadership Grant (2027058). DIG was supported by an NHMRC Senior Principal Research Award (1117766) and subsequently an NHMRC Leadership Grant (2008913). WA was supported by Australian ARC Discovery Early Career Researcher Award (DE220101491) and and Monash FMNHS Future Leader Fellowship. JR is supported by an NHMRC investigator grant (2008981) and an ARC Discovery Project (DP250102065). DPF acknowledges grant support from the ARC Centre of Excellence in Innovations for Peptide and Protein Science (CE200100012), NHMRC Investigator Grant 2009551, and US National Institutes of Health RO1 AI14807-01A1. This research was undertaken in part using the MX2 beamline at the Australian Synchrotron, part of ANSTO, and used the Australian Cancer Research Foundation (ACRF) detector. *Personnel:* We thank staff from the flow cytometry facilities at the Department of Microbiology and Immunology at the Doherty Institute. We thank the staff at the Monash Macromolecular Crystallization Platform for assistance. We would like to thank Conor McNeice for their technical assistance. We thank Professor Andrew Kruse (Harvard Medical School) for providing the naïve yeast nanobody display library, and Edward Harvey (Harvard Medical School) for helpful protocols and discussion for *in vitro* affinity maturation.

## Author contributions

NAG and HFK conceptualised and planned the project. HSH, SJR, WA, CX, CS, LC, APG, HFK and NAG performed the experiments and analysed the data. JYWM, LL, DF, JM, APU, JR and DIG provided key resources and intellectual input. NAG and HFK drafted the manuscript with input from WA and DIG. All authors reviewed the manuscript.

## Data availability

The coordinates of the C11-MR1-5-OP-RU and C11-MR1-Ac-6-FP complex have been deposited in the Protein Data Bank under accession codes 9Q2J and 9Q2K respectively.

