## Supplementary Material for "Specific targeting of MR1-antigen complexes using nanobodies"

**List of Supplementary Materials**

**Fig. S1.** Evolutionary conservation of MR1

**Fig. S2.** Nanobody clone selection

**Fig. S3.** Affinity maturation of clones P1G5 and P2A3

**Fig. S4.** C11 does not block activation of MR1T clones

**Fig. S5.** Characterisation of bispecific antibody species

**Table S1.** Antibodies, dyes and conjugates

**Table S2.** Data collection and refinement statistics.

A.

MR1 α1

```
        1         10        20        30        40        50        60        70        80
Human:  RTHSLRYFRLGVSDPIHGVPEFISVGYVDSHPITTYDSVTRQKEPRAPWMAENLAPDHWERYTQLLRGWQQMFKVELKRLQRHYNHS
Mouse:  ••••••••••A••••GPV•••••••••••••••••••••••••••K•••••••••••••••••••••••••T••A••RH••••••••
```

MR1 α2

```
          90        100       110       120       130       140       150       160       170
Human:  GSHTYQRMIGCELLEDGSTTGFLQYAYDGQDFLIFNKDTLSWLAVDNVAHTIKQAWEANQHELLYQKNWLEEECIAWLKRFLEYGKDTLQRT
Mouse:  •L••••••••••••••••••••••••••••••I•••••••••••M•Y•••IT•••••••L•••Q•••••••••••••••••••••R•••E••
```

Mutations distal to antigen-binding cleft  Mutations proximal to antigen-binding cleft  • conserved with human sequence

B.

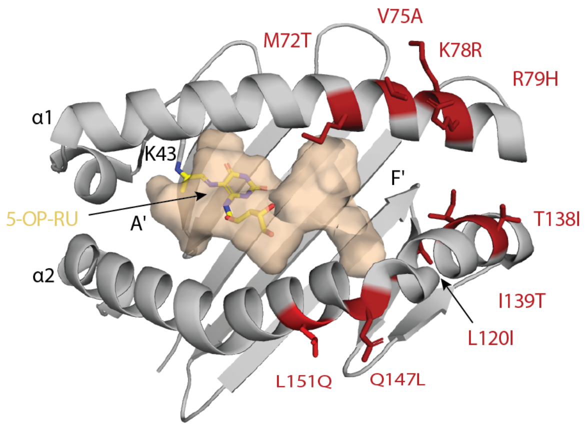


**Figure S1. Evolutionary conservation of MR1. A.** Alignment of amino acid sequences encoding the α1 and α2 domains of MR1 in humans and mice. **B.** Top-down cartoon representation of the crystal structure of human MR1-5-OP-RU (pdb 5D5M) showing the TCR docking platform and antigen-binding cleft with residues highlighted in red that differ between human and mouse that are proximal to the antigen binding cleft or TCR docking platform.

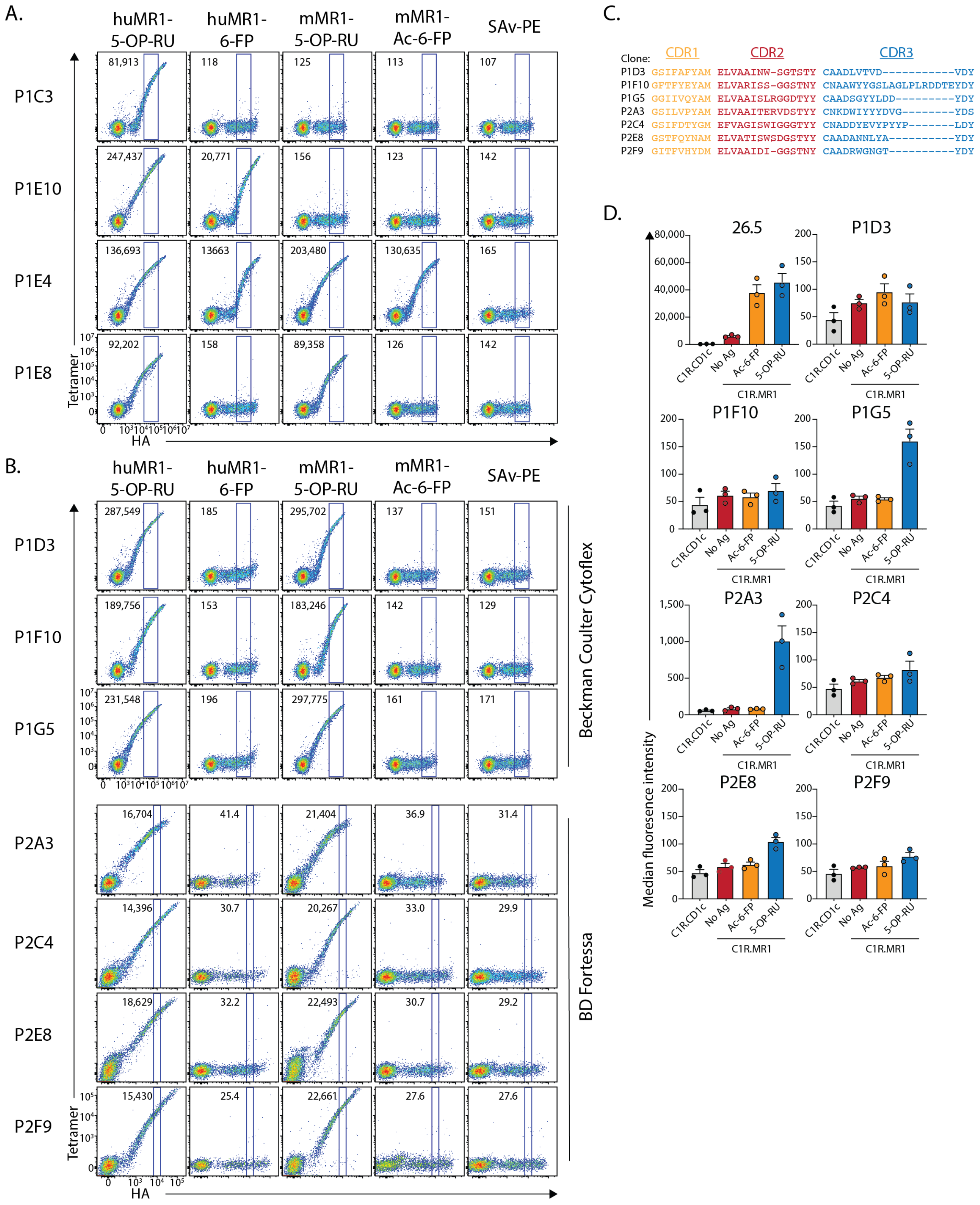


**Figure S2. Nanobody clone selection. A-B.** Flow cytometric pseudocolour plots showing MR1-Antigen tetramer staining on single-cell sorted colonies, demonstrating **A.** the breadth of staining profiles and specificities, and, **B.** those clones with the desired human and mouse MR1-5-OP-RU specificity. **C.** Alignment of CDR amino acid sequences encoding clones in B. **D.** Bar graphs showing median fluorescence intensity of anti-MR1 clone 26.5 antibody or Nb-Fc clones on C1R cells treated with Ac-6-FP or 5-OP-RU. Data points are mean of n=2 technical replicates for n=3 independent experiments. Error bars depict standard error of the mean. Data for 26.5 and P1G5 are the same data that appear in main body figure 2E and are included again here for reference.

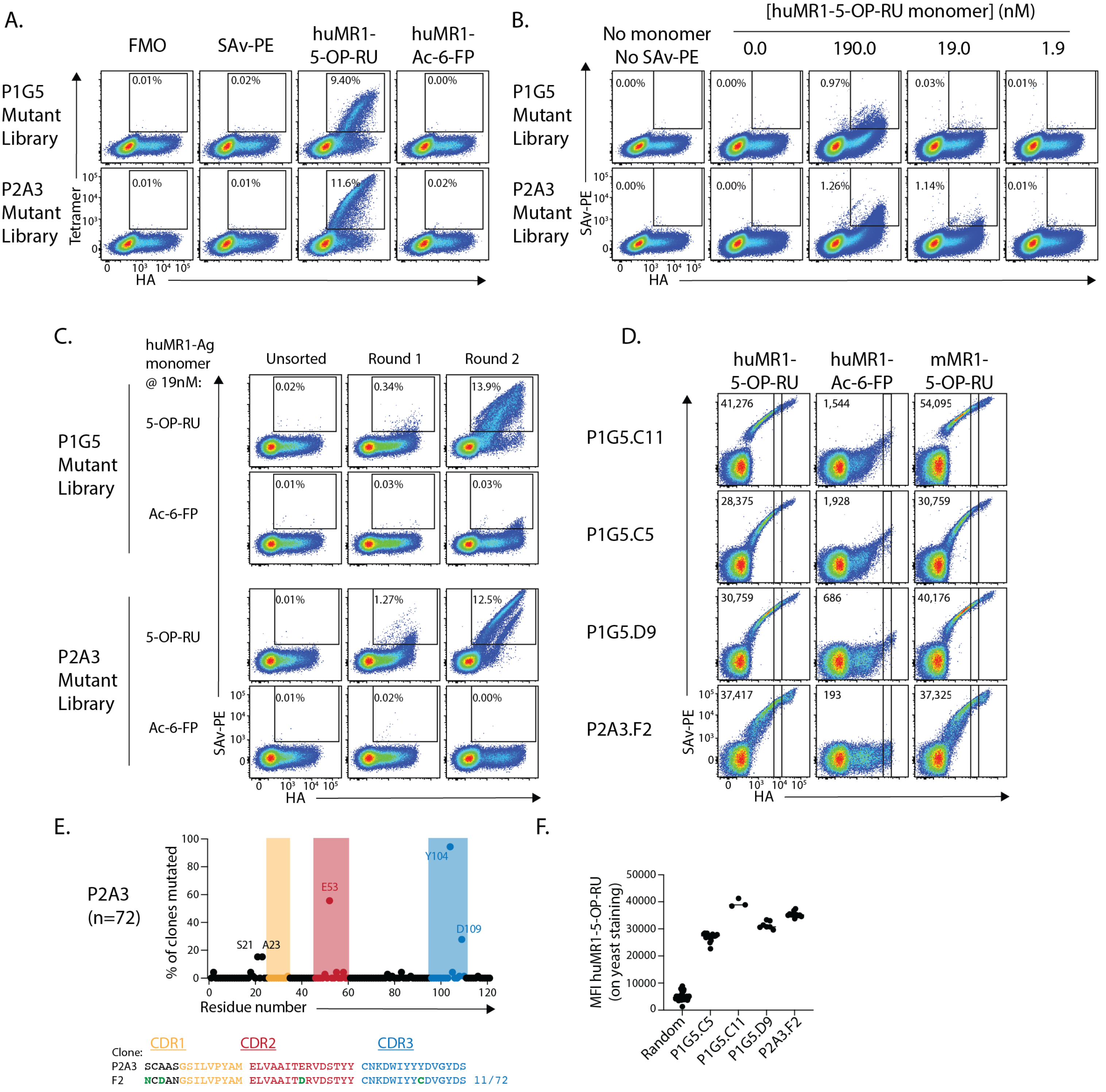


**Figure S3. Affinity maturation of clones P1G5 and P2A3. A-B.** Flow cytometric pseudocolour plots showing MR1-Ag tetramer staining on unsorted yeast mutant libraries stained with **A.** MR1-antigen tetramers or, **B.** varied concentrations of huMR1-5-OP-RU monomers. **C.** Flow cytometric pseudocolour plots showing MR1-antigen monomer staining on P1G5 or P2A3 mutant libraries between subsequent rounds of enrichment. **D.** Flow cytometric pseudocolour plots showing MR1-antigen monomer staining on single-cell sorted colonies from P1G5 or P2A3 mutant libraries with the highest median fluorescence intensities (MFI), depicted numerically on each plot for the gated population. **E.** Scatter plot showing frequency of mutations for individual amino acids of the P2A3 nanobody. Each datapoint represents an individual amino acid and the frequency corresponds to how often that amino acid was mutated across the 72 sequenced clones. **F.** MFI of huMR1-5-OP-RU monomer staining gated as per D for all colonies that had P1G5 C5, D9 or C11 subclones or P2A3 F2 subclone sequences. Colonies with all other sequences are included in the 'random' column. Datapoints for Random, C5, C11 and D9 appear in main body figure 2I and are included here again for reference.

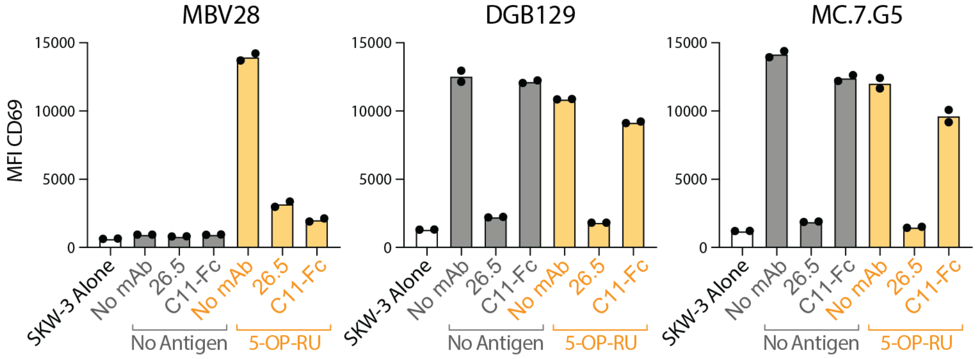


**Figure S4. C11 does not block activation of MR1T clones.** Bar Graph showing median fluorescence intensity (MFI) of CD69 staining on SKW-3.TCR clones MBV28 (control MAIT TCR) or DGB129 and MC.7.G5 (both MR1T clones of distinct specificity) after co-culture with C1R.MR1 cells in the presence or absence of 5-OP-RU and 26.5 anti-MR1 mAb or C11-Fc. Data points represent mean of n=2 technical replicates and data is representative of n=2 independent experiments.

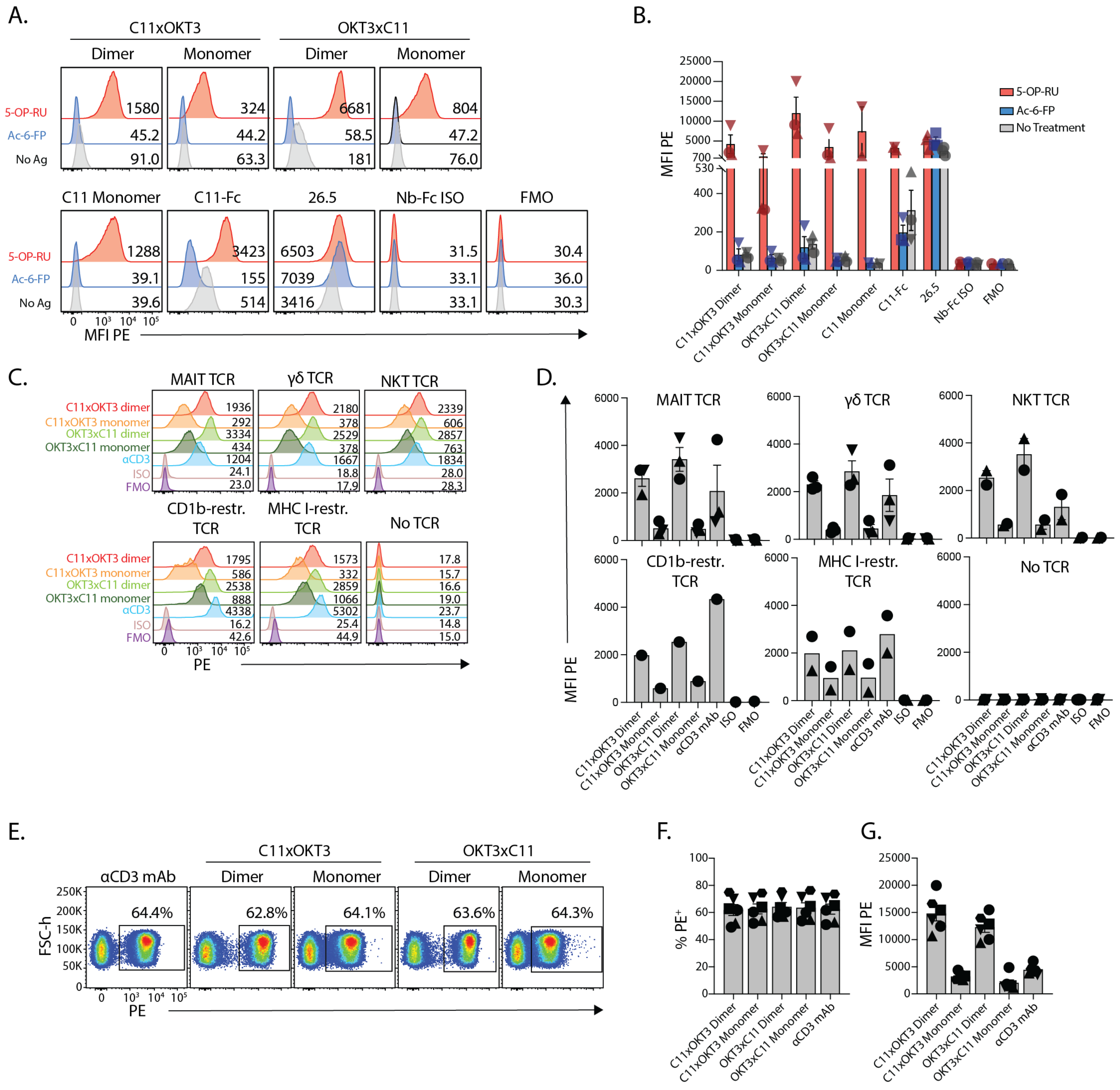

**Figure S5. Characterisation of bispecific antibody species. A.** Representative flow cytometric histogram overlays showing secondary anti-His or anti-Fc staining on C1R.MR1 cells pulsed for 3 hrs with 5-OP-RU at (500 µM), Ac-6-FP (10 µM) or no antigen and subsequently stained with a panel of bispecific antibodies or monomeric nanobody (anti-His) or antibodies (anti-Fc). Numbers next to peaks depict median fluorescence intensity (MFI) of PE secondary antibody staining. **B.** Bar graphs depicting data derived from C from n=3 indpendent experiments. Error bars denote standard error of the mean. **C.** Representative flow cytometric histogram overlays showing secondary anti-His or anti-Fc staining on a panel of SKW-3.TCR cells expressing distinct TCRs stained with a panel of bispecific antibodies or monomeric nanobody (anti-His) or antibodies (anti-Fc). Numbers next to peaks depict MFI of PE secondary antibody staining. **D.** Bar graphs depicting data derived from C from n=3 independent experiments for MAIT, γδ, NKT and no TCR, n=2 for MHC I-restricted TCR and n=1 for CD1b-restricted TCR. Error bars denote standard error of the mean for those with n=3 data points. **E.** Representative flow cytometric pseudocolour plots showing anti-His staining on lymphocytes from PBMC incubated with bispecific antibodies. **F-G.** Bar graphs data from G as (H) the proportion of PE⁺ cells, and (I) the median fluorescence intensity of PE on PE⁺ cells.

**Table S1. Antibodies, dyes and conjugates.**

| Reagent | Fluorochrome | Species Reactivity | Clone | Source |
|---|---|---|---|---|
| Anti-B220 mAb | BV786 | Mouse | RA3-6B2 | Becton Dickinson |
| Anti-CD3 mAb | Unconjugated | Human | OKT3 | BioXCell |
| Anti-CD3 mAb | BUV395 | Human | UCHT-1 | Becton Dickinson |
| Anti-CD4 mAb | BUV395 | Mouse | GK1.5 | Becton Dickinson |
| Anti-CD4 mAb | BUV496 | Human | SK3 | Becton Dickinson |
| Anti-CD8α mAb | BUV805 | Mouse | 53-6.7 | Becton Dickinson |
| Anti-CD8α mAb | BUV805 | Human | SK1 | Becton Dickinson |
| Anti-CD14 mAb | BV570 | Human | M5E2 | BioLegend |
| Anti-CD19 mAb | BV786 | Mouse | 6D5 | Becton Dickinson |
| Anti-CD19 mAb | APC-Cy7 | Human | SJ25C1 | Becton Dickinson |
| Anti-CD19 mAb | BV786 | Human | SJ25C1 | Becton Dickinson |
| Anti-CD25 mAb | BV605 | Mouse | PC61 | BioLegend |
| Anti-CD28 mAb | Unconjugated | Human | 9.3 | BioXCell |
| Anti-CD69 mAb | BUV737 | Mouse | H1.2F3 | Becton Dickinson |
| Anti-CD69 mAb | BUV737 | Human | FN50 | Becton Dickinson |
| Anti-CD69 mAb | PE-Cy7 | Human | FN50 | Becton Dickinson |
| Anti-CD137 mAb | PE | Human | 4B4-1 | BioLegend |
| Anti-CD161 mAb | PE-Cy7 | Human | HP-3G10 | BioLegend |
| Anti-HA tag mAb | AF647 | Hemagglutinin | 16B12 | Biolegend |
| Anti-His tag mAb | PE | N/A | J095G46 | BioLegend |
| Anti-Isotype control IgG1a nanobody | Unconjugated | Mouse | N/A | Made in house |
| Anti-Isotype control mAb IgG2a, k | Unconjugated | Mouse | MOPC-173 | BioLegend |
| Anti-MR1 mAb | PE | Human/Mouse | 26.5 | Biolegend |
| Anti-MR1 mAb | Unconjugated | Human/Mouse | 26.5 | BioLegend |
| Anti-MR1 mAb (for in vivo work) | Unconjugated | Human/Mouse | 26.5 | Purified from hybridoma, provided by J JM (University of Melbourne) |
| Anti-Pan-Immunoglobulin mAb | PE | Mouse | Polyclonal | BioLegend |
| Anti-TCRβ mAb | APC-Cy7 | Mouse | H57-597 | Becton Dickinson |
| Anti-Vα7.2 mAb | BV711 | Human | 3C10 | BioLegend |
| Anti-γδ TCR mAb | PE-Cy7 | Human | 11F2 | Becton Dickinson |
| Fc receptor block | None | Human | N/A | Miltenyi Biotec |
| Fc receptor block | None | Mouse | 2.4G2 | Hybridoma cell line in-house |
| Cell Trace Violet | N/A | Human/Mouse | N/A | Thermo Fisher |
| LIVE/DEAD Fixable Near-IR Dead Cell Stain | Near-IR, Violet | Human/Mouse | N/A | Thermo Fisher |
| 7-AAD Live/Dead | N/A | Human/Mouse | N/A | Sigma |
| Streptavidin-PE | PE | N/A | N/A | Molecular Probes |
| Streptavidin-PE | PE | N/A | N/A | Becton Dickinson |
| Streptavidin-PE-Cy7 | PE-Cy7 | N/A | N/A | Becton Dickinson |

**Table S2. Data collection and refinement statistics.**

| | C11-MR1-5-OP-RU | C11-MR1-Ac-6-FP |
|---|---|---|
| Wavelength | 0.9537 | 0.9537 |
| Resolution range | 47.59 - 2.90 (2.94 - 2.90) | 47.51 - 3.14 (3.18 - 3.14) |
| Space group | P 31 2 1 | P 32 1 2 |
| Unit cell | 208.52 208.52 77.73 90 90 120 | 118.50 118.50 159.06 90 90 120 |
| Total reflections | 290820 (9914) | 141834 (5034) |
| Unique reflections | 83022 (2907) | 43262 (1557) |
| Multiplicity | 3.5 (3.4) | 3.3 (3.2) |
| Completeness (%) | 99.37 (98.04) | 99.00 (92.17) |
| Mean I/sigma(I) | 9.18 (0.91) | 14.26 (0.63) |
| Wilson B-factor | 64.98 | 118.54 |
| R-merge | 0.1149 (1.324) | 0.0556 (1.481) |
| R-meas | 0.1357 (1.571) | 0.06602 (1.766) |
| R-pim | 0.0715 (0.8371) | 0.03487 (0.9415) |
| CC1/2 | 0.995 (0.403) | 0.999 (0.335) |
| CC* | 0.999 (0.758) | 1 (0.708) |
| R-work | 0.1905 (0.3757) | 0.2083 (0.4030) |
| R-free | 0.2044 (0.4134) | 0.2237 (0.3469) |
| Number of non-hydrogen atoms | 4056 | 3727 |
| macromolecules | 3835 | 3692 |
| ligands | 45 | 16 |
| solvent | 176 | 19 |
| Protein residues | 479 | 473 |
| RMS(bonds) | 0.200 | 0.100 |
| RMS(angles) | 4.07 | 2.29 |
| Ramachandran favored (%) | 97.01 | 97.40 |
| Ramachandran allowed (%) | 2.77 | 2.60 |
| Ramachandran outliers (%) | 0.21 | 0.00 |
| Rotamer outliers (%) | 6.33 | 14.13 |
| Average B-factor | 85.30 | 122.47 |
| macromolecules | 86.15 | 122.67 |
| ligands | 61.55 | 99.06 |
| solvent | 72.84 | 103.89 |

Statistics for the highest-resolution shell are shown in parentheses.
